# Molecular Basis for RNA Cytidine Acetylation by NAT10

**DOI:** 10.1101/2024.03.27.587050

**Authors:** Mingyang Zhou, Supuni Thalalla Gamage, Khoa A. Tran, David Bartee, Xuepeng Wei, Boyu Yin, Shelley Berger, Jordan L. Meier, Ronen Marmorstein

## Abstract

Human NAT10 acetylates the N4 position of cytidine in RNA, predominantly on rRNA and tRNA, to facilitate ribosome biogenesis and protein translation. NAT10 has been proposed as a therapeutic target in cancers as well as aging-associated pathologies such as Hutchinson-Gilford Progeria Syndrome (HGPS). The ∼120 kDa NAT10 protein uses its acetyl-CoA-dependent acetyltransferase, ATP-dependent helicase, and RNA binding domains in concert to mediate RNA-specific N4-cytidine acetylation. While the biochemical activity of NAT10 is well known, the molecular basis for catalysis of eukaryotic RNA acetylation remains relatively undefined. To provide molecular insights into the RNA-specific acetylation by NAT10, we determined the single particle cryo-EM structures of *Chaetomium thermophilum* NAT10 (*Ct*NAT10) bound to a bisubstrate cytidine-CoA probe with and without ADP. The structures reveal that NAT10 forms a symmetrical heart-shaped dimer with conserved functional domains surrounding the acetyltransferase active sites harboring the cytidine-CoA probe. Structure-based mutagenesis with analysis of mutants *in vitro* supports the catalytic role of two conserved active site residues (His548 and Tyr549 in *Ct*NAT10), and two basic patches, both proximal and distal to the active site for RNA-specific acetylation. Yeast complementation analyses and senescence assays in human cells also implicates NAT10 catalytic activity in yeast thermoadaptation and cellular senescence. Comparison of the NAT10 structure to protein lysine and N-terminal acetyltransferase enzymes reveals an unusually open active site suggesting that these enzymes have been evolutionarily tailored for RNA recognition and cytidine-specific acetylation.

## INTRODUCTION

Nature uses chemical modifications to diversify the functional properties of proteins and nucleic acids. One prototypical example of this is acetylation. In mammalian cells, the vast majority of proteins are acetylated on the epsilon amine of lysine ^1,2^ or at their N-terminus ^3-5^. These modifications are catalyzed by enzymes exhibiting homology to the GCN5-related N-acetyltransferase (GNAT) family ^6^. Nucleic acid acetylation also exists, and in humans occurs in the form of N4-acetylcytidine (ac^4^C) in RNA. In contrast to protein acetylation - which is dependent on several enzyme activities ^3,5,7,8^ - cytidine acetylation is catalyzed by a single GNAT-containing protein, *N*-acetyltransferase 10 (NAT10) ^9-13^. The importance of ac^4^C is implied by the fact that it occurs across all domains of life and that NAT10 is essential in all known eukaryotes, ranging from vertebrates to yeast ^9,12,14,15^.

The domain architecture of NAT10 is distinct from other mammalian acetyltransferase enzymes. In addition to the aforementioned GNAT fold ^9,14,16^, NAT10 also possesses an N-terminal ATP-dependent helicase domain, a C-terminal RNA binding domain, and several other regions of notable sequence homology, resulting in a relatively large polypeptide chain of ∼120 kDa ^16,17^. Although initially reported to be a histone and tubulin acetyltransferase ^18,19^, more recent studies strongly suggest that NAT10 is a dedicated RNA N4-cytidine acetyltransferase, with a role in RNA biogenesis through the acetylation of specific cytidine targets on 18S rRNA (C1280/C1337 on helix 34, and C1773/C1842 on helix 45 in *S. cerevisiae* and H. *sapiens* respectively) and tRNA (C12 in eukaryotic serine and leucine tRNA) ^9,11,13,14^. Context-dependent roles in enhancing translation efficiency through mRNA acetylation have also been proposed ^10,20^. A NAT10 adaptor protein, Thumpd1 (Tan1 in yeast), has been shown to facilitate tRNA acetylation ^9,21^, while two box C/D small nucleolar RNAs (snoRNAs; snR4 and snR45 in yeast) have been demonstrated to help guide the yeast NAT10 homologue (Kre33) to its rRNA targets (snR4 for C1280 on helix 34 and snR45 for C1773 on helix 45) ^22^. SNORD13 plays a similar role in guiding acetylation to C1842 on helix 45 in human rRNA ^23-25^. In addition to its catalytic acetyltransferase activity, NAT10 appears to contribute to the proper assembly and maturation of the 90S pre-ribosome via non-catalytic roles, including by making interactions with other ribosome biogenesis factors ^26-28^.

Although it is an essential protein, data indicates NAT10 may be a viable therapeutic target for the treatment of various cancers and cellular aging, and has been implicated in the regulation of p53 ^17,29-35^, as well as the human accelerated aging genetic disorder, Hutchinson-Gilford Progeria Syndrome (HGPS) ^36,37^. Heterozygous knockout of NAT10 has been demonstrated to modulate HGPS phenotypes in a mouse model ^38^. These phenotypes can be recapitulated with the treatment of a putative NAT10 inhibitor; however, the basis for this is unclear as this molecule was later found to possess pleiotropic off-target activity and not bind NAT10 or inhibit RNA acetylation ^39^. Overall, genetic and biological analyses highlight the importance of NAT10 in aging pathways and disease, as well as the importance of research that could facilitate the development of more drug-like chemical probes of RNA cytidine acetylation.

Structural studies of a bacterial acetyltransferase have yielded valuable information about the molecular mechanism of RNA acetylation. A crystal structure of TmcA, a bacterial *S. enterica* homolog of NAT10 that shares common functional domains, was determined in complex with acetyl-CoA and ADP ^40^. TmcA reveals a monomeric L-shaped morphology, with the helicase domain together with a structured DUF699 domain mimicking DEAD-box RNA helicases lying against the GNAT domain, which is followed by the tail domain that is proposed to contribute to tRNA binding ^14^. Structural analyses of eukaryotic NAT10 has been limited to cryo-EM structures of the 90S pre-ribosome from *Chaetomium thermophilum* (*Ct*) and *Saccharomyces cerevisiae* (*Sc*), where it was found that NAT10 (fungal Kre33) forms a head-to-head and tail-to-tail homodimer on the ribosome and cooperates with Bms1, Enp2, Brf2 and Lcp5 proteins to mediate the final compaction of 18S rRNA subdomain onto the 90S pre-ribosome ^41,42^.

Despite this progress, the molecular features of NAT10 responsible for its cytidine-specific RNA acetylation activity remain unresolved. A detailed mechanistic understanding of NAT10 could enable a better understanding of the biological activity of this enzyme as well as setting the stage for chemical probe development. For these reasons, we set out to determine the structure of *Chaetomium thermophilum* NAT10 (*Ct*NAT10) bound to a cytidine-CoA probe with and without ATP, and carried out associated functional studies *in vitro* and in yeast and mammalian cells. These studies reveal that NAT10 is a symmetrical heart-shaped dimer with conserved functional domains from both subunits surrounding two acetyltransferase active sites. Structure-based mutagenesis indicates a critical role for two conserved active site residues and two basic patches proximal and distal to the active site for RNA-specific acetylation. Applying these insights in functional studies, we corroborate the importance of NAT10 catalytic activity in yeast fitness under temperature stress and demonstrate this property can also impact cellular senescence in mammalian cells. Finally, we compare NAT10 to histone and N-terminal acetyltransferase structures, which reveals a more open active site that appears more tailored for RNA-specific acetylation. Overall, our studies provide molecular level insights into eukaryotic RNA acetyltransferase activity, providing a foundation for domain-selective inhibition and functional studies of NAT10 in human disease.

## RESULTS

### Recombinant NAT10 is active for RNA acetylation and binding and can be bound by a cytidine-CoA conjugate probe

To facilitate biochemical and structural studies of NAT10, we first expressed recombinant full length *Chaetomium thermophilum* NAT10 (*Ct*NAT10) from both *Escherichia coli* (*E.coli*) and *Spodoptera frugiperda* (*Sf9*) (**Supplemental Figure 1**). NAT10 in this thermophilic fungus contains functional domains that are conserved from bacteria to human, with a notably higher degree of evolutionary conservation of sequence in human. (**Supplemental Figure 2**). To evaluate the catalytic activity of recombinant *Ct*NAT10, we carried out an autoradiography activity assay against a rRNA substrate h45 (chemically synthsized 18S rRNA helix 45), which revealed increasing activity over time with saturation (**Figure 1A**). Next, we evaluated the RNA binding activity of *Ct*NAT10 by carrying out a gel shift assay utilizing a series of RNA substrates. This assay demonstrated concentration-dependent RNA binding activity by the recombinant protein, consistent with its known function (**Figure 1B**). To assess the ability of *Ct*NAT10 to bind to cofactors, we applied a differential scanning fluorimetry (DSF) assay (**Figure 1C**). While apo *Ct*NAT10 displayed a 42.0 ± 0.3 °C melting temperature (T_m_), we observed statistically significant thermal stabilization in the presence of Mg^2+^ - ATP (53.8 ± 0.1 °C), acetyl-CoA (46.1 ± 0.1 °C) and their non-hydrolysable analogs. Surprisingly, binding of the tRNA and rRNA substrates did not significantly increase *Ct*NAT10 thermostability, suggesting that these molecules may bind predominantly to the *Ct*NAT10 surface.

**Figure 1.**
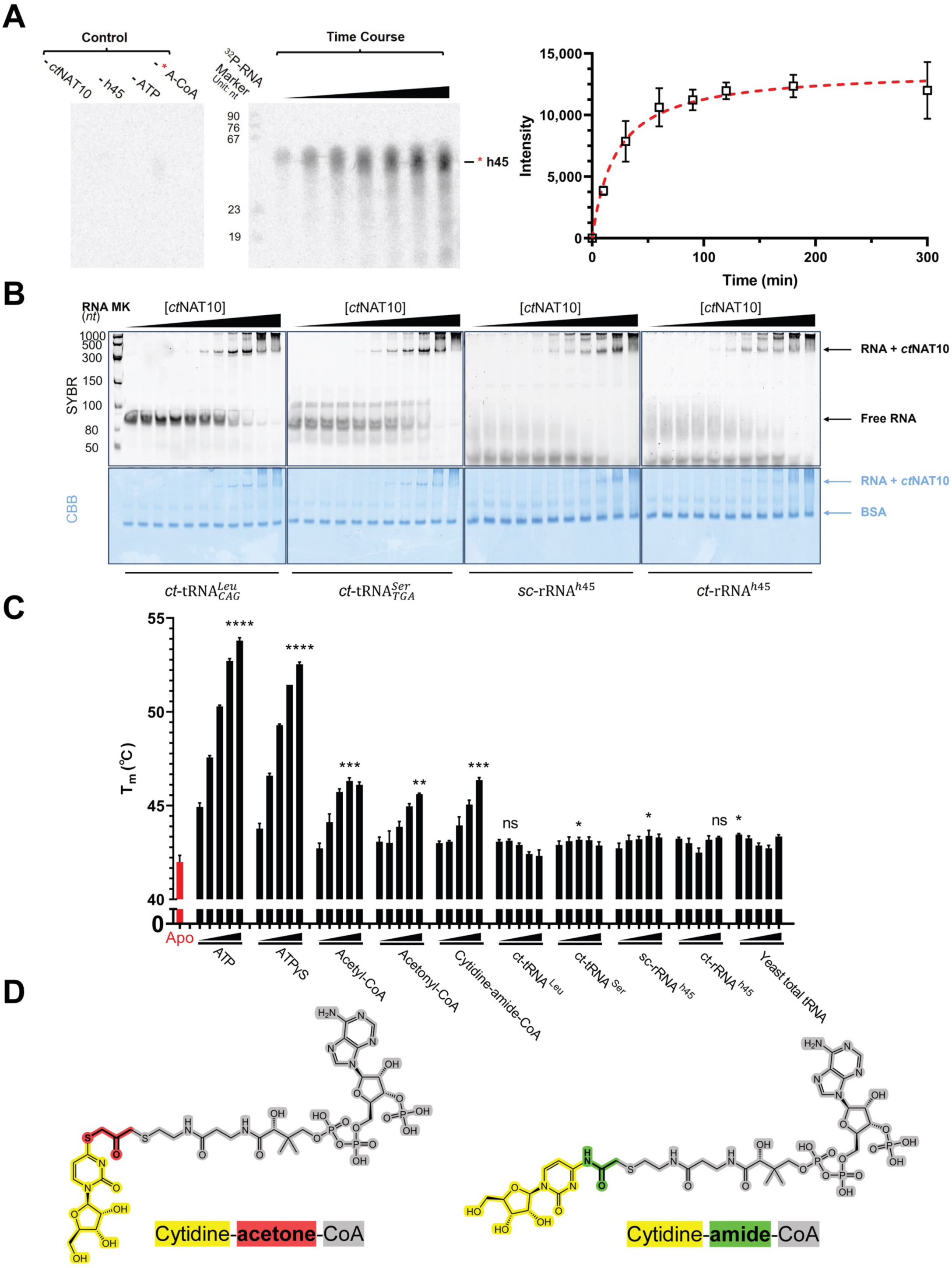
Biochemical characterization of recombinant NAT10 and cytidine-CoA probe compounds. **(A)** Left **-** Autoradiography of *Ct*NAT10 catalyzed *in vitro* acetylation of a h45 RNA substrate as a function of component deletion (60 min.) and time (10, 30, 60, 90, 120, 180, 300 min.). The level of ^14^C labeled h45 (* h45) was separated by 15% denaturing urea polyacrylamide gel electrophoresis, and detected by phosphor screen exposure. Right - Plot of the data derived from gel image. Averaged values of three independent experiments with S.D. values are shown. **(B)** Electrophoretic mobility shift assays (EMSAs) of purified *Ct*NAT10 wild-type binding with synthesized tRNAs and rRNAs. 6% polyacrylamide native gels were stained with SYBR™ Gold (upper panel) and Coomassie Brilliant Blue (lower panel) to visualize RNA and protein, respectively. RNA input was fixed at 5 pmol and *Ct*NAT10 was titrated at 0, 0.2, 0.4, 0.7, 1.5, 3, 6, 11, 23, 45, 90 pmol (from left to right). 0.2 mg/ml BSA was added to each sample as a loading control and to prevent non-specific binding. RNA-*Ct*NAT10 complex bands, free RNA probe bands and BSA bands are indicated with arrows. **(C)** Differential Scanning Fluorimetry (DSF) evaluation of *Ct*NAT10 (1 μM) in the absence (red bar) and presence of substrates, metabolic ligands and their non-hydrolysable analogs, and bisubstrate cytidine-CoA probe (black bars). Each ligand was evaluated from low to high concentration as follows (ATP: 0.25, 2.5, 25, 250, 2500 μM; ATPɣS: 0.25, 2.5, 25, 250, 2500 μM; Acetyl-CoA: 0.25, 2.5, 25, 250, 2500 μM; acetonyl-CoA: 0.25, 2.5, 25, 250, 2500 μM; Cytidine-amide-CoA: 0.20, 2.0, 20, 200, 2000 μM; ct-tRNA^Leu^: 0.625, 1.25, 2.5, 5, 10 μM; ct-tRNA^Ser^: 0.625, 1.25, 2.5, 5, 10 μM; sc-rRNA^h45^: 0.625, 1.25, 2.5, 5, 10 μM; ct-rRNA^h45^: 0.625, 1.25, 2.5, 5, 10 μM; Yeast total RNA: 2.5, 5, 10, 20, 40 μM). Averaged values of three technical replicates with S.D. values are shown for each condition. P-values were calculated only for the melting temperature of each ligand at the concentration that maximizes the thermal stability of *Ct*NAT10 relative to the apo protein. All the P-values were calculated by Brown-Forsythe and Welch ANOVA with Dunnett’s T3 multiple comparisons test, with individual variances computed for each comparison (ns: not significant, ∗: P<0.05, ∗∗: P<0.01, ∗∗∗: P<0.001, ∗∗∗∗: P<0.0001)**. (D)** The sub-structure portions of cytidine-CoA bisubstrate probes are highlighted (cytidine in yellow, bridging linkers in red and green, and CoA in grey).

Bisubstrate probes of acetyltransferases have proven valuable tools for studying the structure and function of this enzyme class ^43-45^. Therefore, we synthesized putative bisubstrate cytidine-CoA conjugate probes designed to mimic the NAT10 reaction intermediate and facilitate structural studies. Two compounds were synthesized, cytidine-acetone-CoA and cytidine-amide-CoA, whose structures differ only in the presence of an acetone or amide linker, respectively (**Figure 1D**). The cytidine-amide-CoA probe (46.4 ± 0.1 °C) demonstrated thermal stabilization comparable to acetyl-CoA, suggesting a similar mode of interaction and a minimal contribution of the cytidine to binding energy (**Figure 1C**).

### NAT10 forms a symmetrical heart-shaped dimer with domains from both subunits surrounding the acetyltransferase active site

Having demonstrated the production of active recombinant *Ct*NAT10, we next set out to prepare liganded protein for single particle cryo-EM structure determination. We were able to produce three structures: *E.coli* produced recombinant *Ct*NAT10/cytidine-acetone-CoA, *E.coli* produced recombinant *Ct*NAT10/cytidine-amide-CoA/ADP and *Sf9* produced recombinant *Ct*NAT10/cytidine-amide-CoA/ADP at overall resolutions of 3.3 Å, 3.2 Å, and 3.0 Å, respectively (**Table 1, Supplemental Figures 3, 4 and 5)**. The *Ct*NAT10/cytidine-acetone-CoA structure was initially modeled first using *E. coli* TmcA (*Ec*TmcA) as a template. As the micrographs suggested two-fold symmetry, C_2_ symmetry was applied during refinement **(Supplemental Figure 3)**. The *Ct*NAT10/cytidine-acetone-CoA model was then used to build both the *E.coli* and *Sf9* produced *Ct*NAT10/cytidine-amide-CoA/ADP models. The three structures superimpose well showing no significant conformational changes due to the presence of ADP or cytidine-CoA probes with an RMSD between all Cα atoms of less than 1.30 Å (**Supplemental Figure 6**). All the previously reported functional domains are well-resolved in the three structures analyzed and the regions corresponding to these functional domains exhibit minor variations from the residue ranges documented in prior reports (**Figure 2A**). Additionally, a novel N(o)LS domain was defined, which contains a putative nuclear localization signal (NLS) and nucleolar localization signal (NoLS) motifs that only exist in eukaryotic NAT10 proteins ^16^. Each subunit of the two-fold symmetric *Ct*NAT10 dimer contains 31 helices and 22 strands forming three sheets within the N(o)LS/DUF1726, helicase and GNAT domains while the RNA binding domain is exclusively helical (**Figure 2B and Supplemental Figure 7**).

**Figure 2.**
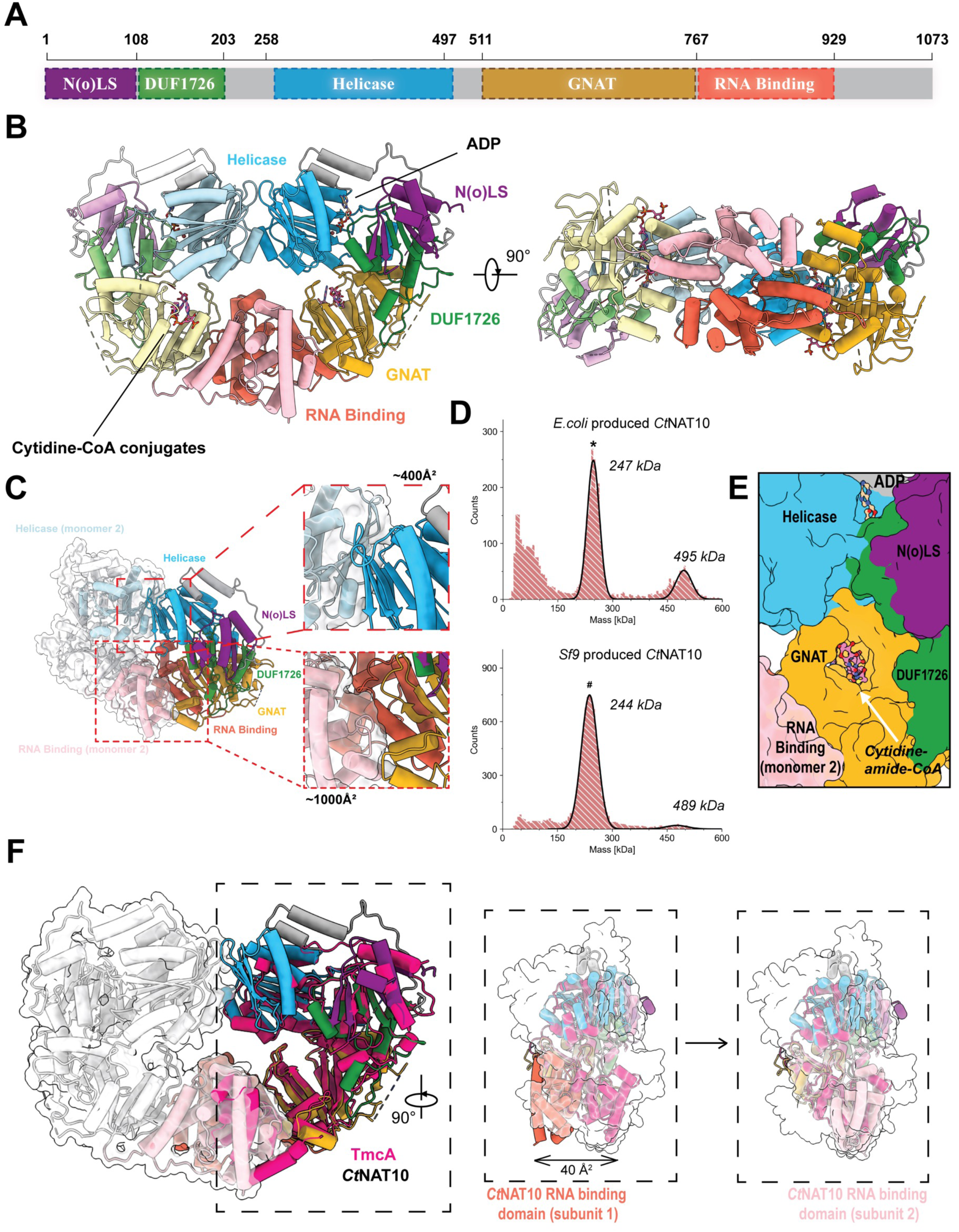
Overall structure of *Ct*NAT10 and comparison to TmcA. **(A)** Scaled schematic representation of *Ct*NAT10 domain organization, with nomenclature and residue count. **(B)** Color-coded cartoon of overall *Ct*NAT10 dimer structure (subunits shown in light and dark shade of color) using the *Sf9* produced recombinant *Ct*NAT10/cytidine-amide-CoA/ADP structure as the representative. Each domain is colored and named corresponding to the domain schematic representation in A. **(C)** Structural features associated with *Ct*NAT10 dimerization. One subunit of the *Ct*NAT10 dimer is colored like B, while the domains of the other subunit are depicted as a ghostly white cartoon and surface representation, with the exception of the RNA binding (light salmon) and helicase (light sky blue) domains that directly participate in dimerization. **(D)** Characterization of the oligomeric state of recombinant *Ct*NAT10 by mass photometry. Histograms in panels show the particle counts of *E.coli* (up) and *Sf9* (bottom) produced recombinant *Ct*NAT10 at the indicated molecular mass. Black lines are Gaussian fits to the peaks and the mass of the Gaussian fit are indicated above each peak. In both samples, *Ct*NAT10 was mostly present as a dimer (theoretical MW of *Ct*NAT10 dimer: 241.02 kDa), and the dimer peaks for *E.coli* (*) and *Sf9* (^#^) sample are indicated. **(E)** Close-up view of the GNAT active site. Each domain is colored and named corresponding to the domain schematic representation in A. ADP and cytidine-amide-CoA are depicted in ball-and-stick representation. **(F)** Superimposition of *Ct*NAT10 and *E. coli* TmcA. TmcA is colored in hot pink. *Ct*NAT10 subunit 1 is colored as in A while subunit 2 is depicted in ghostly white with the exception of the RNA binding domain (light salmon). The insert is rotated 90° with the RNA binding domain of TmcA superimposed with the *Ct*NAT10 RNA binding domains of subunits 1 (left) and 2 (right).

**Figure 3.**
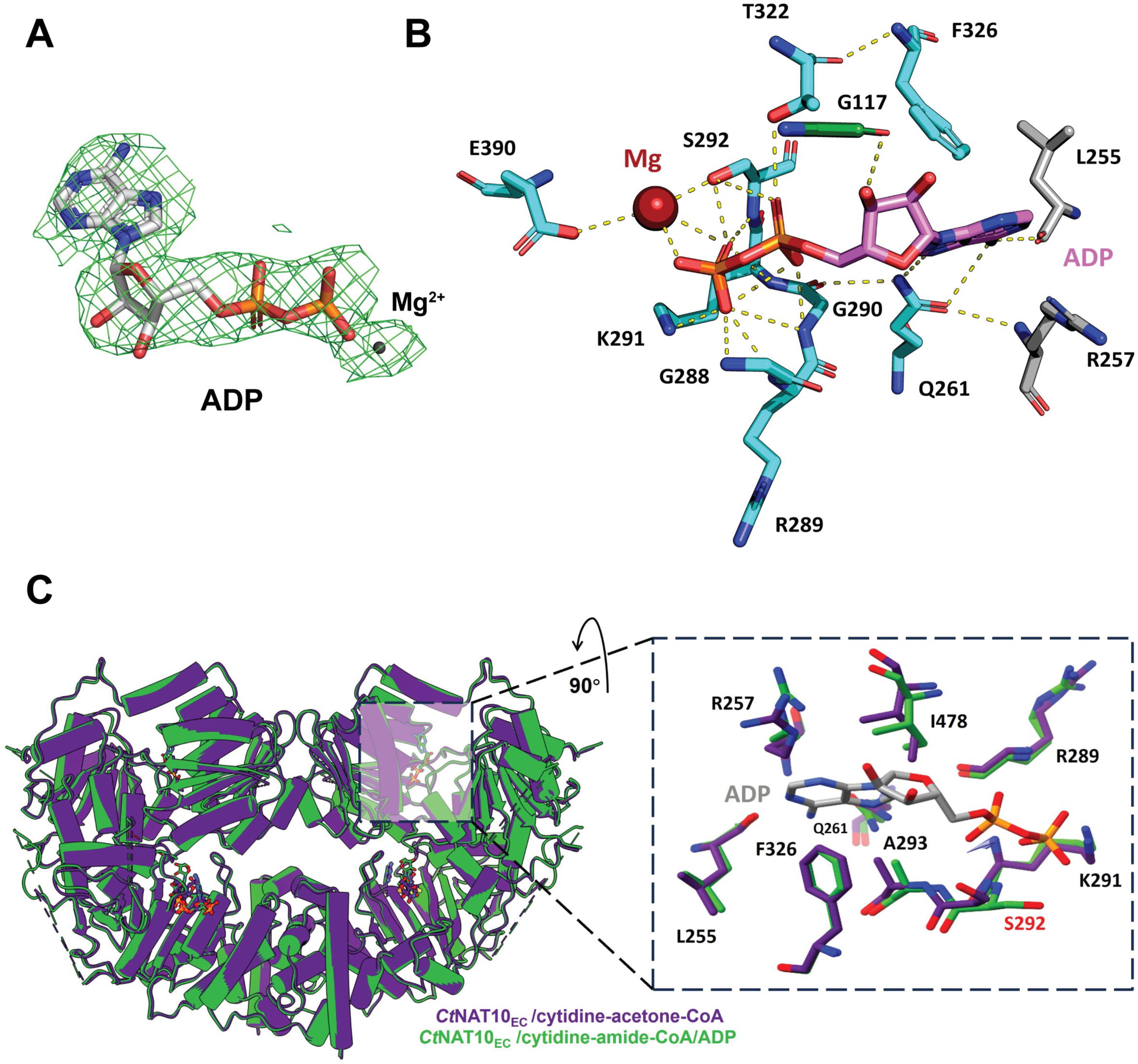
ATP binding site of *Ct*NAT10. **(A)** Cryo-EM density of ADP and Mg^2+^ ion. A contour level of 8 σ was applied. **(B)** Close-up view of ADP/Mg^2+^ - protein interactions. Bound ADP (magenta) and key residues involved in interactions with ADP/Mg^2+^ are drawn as sticks and are labelled. The Mg^2+^ ion (maroon) is shown as a spherical ball. Hydrogen bonds are indicated by yellow-colored dashed lines, with pi-pi interaction and hydrophobic interaction not explicitly shown. Atoms are colored as follows: C -the same as the domain color where the residue is located, N -blue, S - orange and O -red. **(C)** Left - Structure comparison of *E.coli* produced *Ct*NAT10 bound to bisubstrate cytidine-CoA probe analog with and without ADP. *Ct*NAT10/cytidine-acetone-CoA is shown in purple and *Ct*NAT10/cytidine-amide-CoA/ADP is shown in green. Right - Selected residues involved in interactions with ADP are drawn as sticks and are labelled in the zoom-in view and shown with the same color-coding as in the left. Ser292, the only residue that changes orientation to accommodate the binding of ADP, is highlighted with a red label.

**Figure 4.**
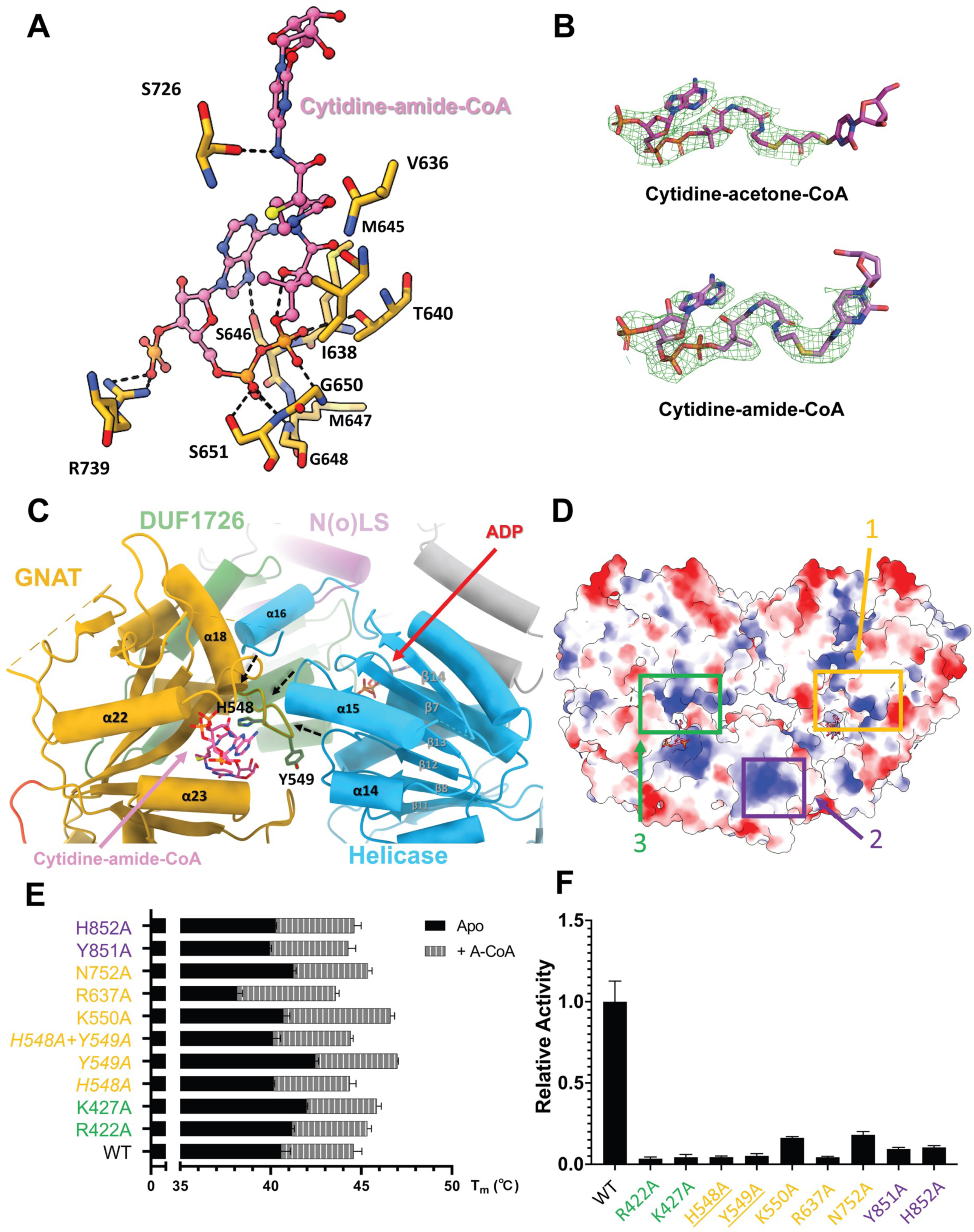
Acetyl-CoA, Cytidine and RNA binding sites of *Ct*NAT10. **(A)** Cryo-EM density of Cytidine-acetone-CoA and Cytidine-amide-CoA. Contour levels of 7 σ and 8 σ was shown, respectively. **(B)** Close-up view of interactions between Cytidine-amide-CoA and *Ct*NAT10 protein. Cytidine-amide-CoA (pink) is drawn with ball-and-stick representation and labeled with the same color, and residues that make hydrogen bond and with Cytidine-amide-CoA are shown by black-colored dashed lines. Van der Waals interaction and hydrophobic interaction are not explicitly shown but described in the main text. Atoms are color-coded as in Figure 3. **(C)** Putative catalytic residues His548 and Tyr549 (green) within the GNAT domain (gold) are shown proximal to the helicase domain (green) where movement of the helicase domain can influence positioning of these residues. Bound ADP (red) and Cytidine-amide-CoA (lavender) are drawn as sticks and are labelled with corresponding color. Dotted black arrows are drawn to suggest how the helicase domain may modulate the positioning of His548 and Tyr549. **(D)** Electrostatic surface diagram of the *Ct*NAT10 structure, highlighting 3 color-coded regions (yellow, purple, and green) selected for mutagenesis and functional studies. Neutral, electropositive and electronegative surfaces of the *Ct*NAT10 structure are colored white, blue and red, respectively. **(E)** DSF evaluation of *Ct*NAT10 mutant constructs in the absence (solid black bar) and presence (patterned bar) of acetyl-CoA (A-CoA). The residues are color-coded according to the regions in D. Averaged values of four technical replicates with S.D. values are shown for each dataset. **(F)** Relative enzyme activity of *Ct*NAT10 mutant constructs relative to wild type (WT). Residues are color-coded according to the regions in D. Putative catalytic residues (His548 and Tyr549) are underlined. Data presented as mean ± standard deviation, n= 3 technical replicates. Plot of the data derived from gel image (see “Supplemental figure 8C” upper panel). The data were calculated from the autoradiogram band intensities.

**Figure 5.**
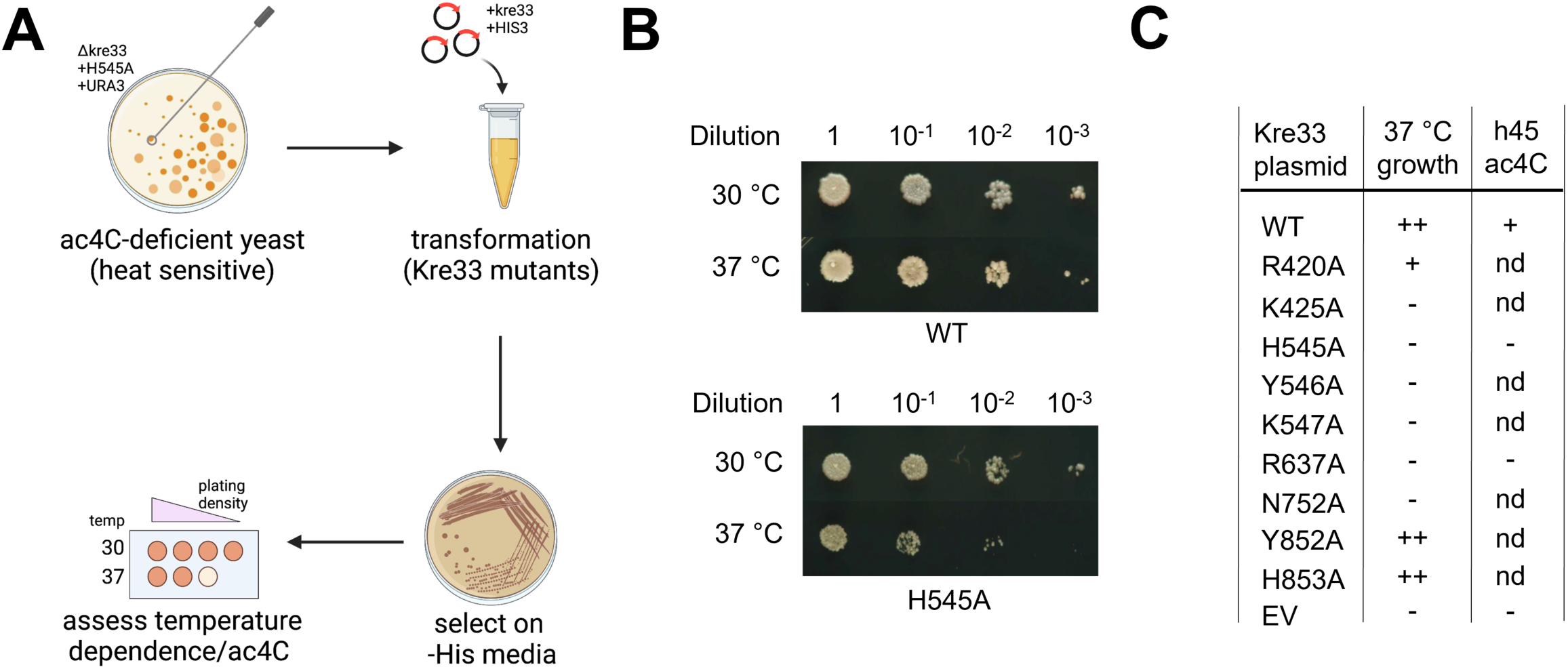
*In vivo* assessment of conserved NAT10 residues by functional complementation. **(A)** Scheme for yeast functional complementation assay. Assays were performed in a parent yeast strain (Δkre33 + pSH35) in which the endogenous kre33 allele has been deleted replaced with a *Sc*Kre33 H545A (*Sc*H545A, corresponding to *Ct*H548A) variant. The lack of Kre33 catalytic activity causes this strain to exhibit a growth defect at high temperatures ^9^. Yeast were transformed individually with kre33 plasmids expressing the indicated mutants under TDH3 control with a His3 selection marker. Transformants were grown on -His media and transformation confirmed by PCR amplification and sequencing. Serial 10-fold dilutions were plated on YPD media and incubated for 2 d at the indicated temperatures to assess rescue of temperature-sensitive growth. **(B)** Validation of assay demonstrating that temperature sensitive growth is rescued by WT *Sc*Kre33 (top) but not *Sc*H545A mutant (bottom). Strains were grown, diluted, plated on YPD plates, and grown at the indicated temperatures as described in A. **(C)** Summary of rescue of temperature-sensitive growth defects and RNA acetylation by Kre33 mutants. ++, complete rescue of growth defect; +, partial rescue of growth defect or RNA acetylation; -, no rescue of growth defect or RNA acetylation. nd = not determined. Growth and RNA acetylation assay data is provided in the Supplementary Information.

**Table 1.** Cryo-EM data collection, refinement, and validation statistics.

|  | CtNAT10<sub>EC</sub> /cytidine-acetone-CoA<br>EMD-43996<br>PDB: 9AYM | CtNAT10<sub>EC</sub> /cytidine-amide-CoA/ADP<br>EMD-44038<br>PDB: 9B0E | CtNAT10<sub>Sf9</sub> /cytidine-amide-CoA/ADP<br>EMD-44042<br>PDB: 9B0I |
| --- | --- | --- | --- |
| **Data collection and processing** |  |  |  |
| Magnification | 105,000 | 105,000 | 105,000 |
| Voltage (keV) | 300 | 300 | 300 |
| Electron exposure (e/Å<sup>2</sup>) | 45 | 45 | 42.9 |
| Defocus range (μm) | -1.0 to -2.5 | -1.0 to -3.0 | -1.0 to -3.0 |
| Pixel size (Å) | 0.415 | 0.415 | 0.43 |
| Number of images | 5,500 | 7,351 | 5,986 |
| Symmetry imposed | C2 | C2 | C2 |
| Initial particles (no.) | 1,228,342 | 407,808 | 5,219,352 |
| Final particles (no.) | 160,385 | 73,181 | 162,769 |
| Map resolution (Å) | 3.29 | 3.19 | 3.02 |
| FSC threshold | 0.143 | 0.143 | 0.143 |
| **Refinement** |  |  |  |
| Initial model used (PDB code) | 2ZPA | 2ZPA | 2ZPA |
| Model resolution (Å) | 3.4 | 3.4 | 3.2 |
| FSC threshold | 0.5 | 0.5 | 0.5 |
| Map sharpening *B* factor (Å<sup>2</sup>) | -151.2 | -109.9 | -128.6 |
| **Model composition** |  |  |  |
| Non-hydrogen atoms | 12,950 | 13,158 | 13,040 |
| Protein residues | 1,624 | 1,642 | 1,628 |
| Ligands | 2 | 4 | 4 |
| ***B* factors (Å<sup>2</sup>)** |  |  |  |
| Protein | 71.02 | 112.55 | 45.32 |
| Ligand | 69.68 | 107.17 | 37.18 |
| **R.M.S. deviations** |  |  |  |
| Bonds lengths (Å) | 0.005 | 0.002 | 0.003 |
| Bond angles (°) | 0.678 | 0.642 | 0.624 |
| **Validation** |  |  |  |
| MolProbity score | 1.58 | 1.41 | 1.32 |
| Clash score | 10.09 | 7.55 | 4.21 |
| Poor rotamers (%) | 0.28 | 0.28 | 0.14 |
| **Ramachandran plot** |  |  |  |
| Favored (%) | 97.76 | 98.21 | 97.39 |
| Allowed (%) | 2.24 | 1.79 | 2.61 |
| Disallowed (%) | 0 | 0 | 0 |

Unlike *Ec*TmcA, which is an L-shaped monomer, the *Ct*NAT10 is a head-to-head, and tail-to-tail heart-shaped dimer that is stabilized by head domain interactions between helicase domains and tail domain interactions between the RNA binding domains that buries solvent excluded surfaces of ∼400 Å^2^ and ∼1000 Å^2^, respectively (**Figure 2C**). Notably, the residues involved in dimerization are ∼90% conserved through eukaryotic NAT10 proteins, but not in *Ec*TmcA (**Supplemental Figure 2**). Mass photometry of the *Ct*NAT10 at a concentration of 10 nM also shows the presence of a *Ct*NAT10 dimer, together supporting the biological relevance of a *Ct*NAT10 dimer (**Figure 2D**), even in the absence of ribosome binding ^41^. Importantly, the *Ct*NAT10 dimer places each of the functional domains around the symmetric GNAT active sites, with the RNA binding domain coming from the symmetry related subunit (**Figure 2E**).

Although *Ct*NAT10 and *Ec*TmcA share only 24.8% sequence identity (calculated by Clustal Omega ^46^), all the functional domains align well (RMSD of 2.471 Å over 301 common Cα atoms), with the exception of the RNA binding domain (**Figure 2F**). Compared to *Ec*TmcA, in the *Ct*NAT10 structure, the RNA binding domain shifts ∼40 Å towards the symmetry related *Ct*NAT10 subunit. Interestingly, the RNA binding domain of the symmetry related subunit superimposes much more closely to the position of the RNA binding domain of *Ec*TmcA (**Figure 2F**), thus further supporting the biological importance of the dimeric state of NAT10.

### ADP binds at the helicase-DUF domain interface and does not significantly alter the overall *Ct*NAT10 structure

While ATP was added to the cryo-EM samples with bound nucleotide, only ADP could be modeled into the cryo-EM density, suggesting that *Ct*NAT10 had catalyzed ATP hydrolysis in the absence of a cognate cytidine substrate. ADP and a bound Mg^2+^ ion are well resolved in the *Sf9* produced recombinant *Ct*NAT10/cytidine-amide-CoA /ADP structure (**Figure 3A**). The observation that added ATP could only be modeled as ADP was also made in the structure of *Ec*TmcA ^40^, where the residues that are involved in ADP coordination are largely conserved (**Figure 3B and Supplemental Figure 2**). In the *Ct*NAT10/cytidine-amide-CoA/ADP structure, the ADP and Mg^2+^ are wedged in between the helicase and DUF domains, and most of the protein interactions are mediated by the α-helix bundle (α9-α12) from the helicase domain (**Supplemental Figure 7**). The adenosine group makes hydrophobic interactions with either the side chain or main chain carbon atoms of residues Gly117, Arg257, Gln261, pi-pi interaction with Phe326, and hydrogen bond interactions with residues Leu255, and Gln261. The pyrophosphate moiety of ADP is mainly held by the side chain of Lys291, several backbone amide nitrogen atoms spanning residues Gly288-Ser292, and two hydrogen bonding interactions from Thr322 with the α-phosphate and the side chain of Ser292 with the β-phosphate. Mg^2+^ is chelated by the β-phosphate of ADP and the side chain of Ser292 and Glu390. A comparison of the *Ct*NAT10/cytidine-CoA probe structures with and without ADP reveals that ADP binding does not make significant changes to the overall structure of *Ct*NAT10, with an RMSD of 0.506 Å over 1,479 common Cα atoms (**Figure 3C and Supplemental Figure 6**). The only residue that changes its orientation upon ADP binding is Ser292, which rotates by ∼180° **(Figure 3C)**. Together, these findings demonstrate that ATP binding and hydrolysis does not significantly change the NAT10 structure, at least in the absence of a cognate cytidine containing RNA substrate.

### The active site tunnel is surrounded by conserved domains and implicates important residues for catalysis

As expected, the cytidine-CoA probes are bound to the GNAT domain of *Ct*NAT10, which is located roughly in the middle of each of the protein subunits. In the *Ct*NAT10/cytidine-amide-CoA/ADP structure, the adenosine and pyrophosphate group of the cytidine-CoA probe is mainly clamped into place by van der Waals interactions and hydrogen bonds between Met647, the backbone amides of residues Ser646, Gly648, Gly650 and the side chains of residues Thr640, Ser646, Ser651, with the pyrophosphate group; and the side chains of residues Ser646 and Arg739 with the adenine base and 3′-phosphate group of the adenosine. The pantetheine arm rests inside a largely hydrophobic tunnel and stabilized by several hydrophobic interactions from Ile638 and Met645 (**Figure 4A**). Consistent with most GNAT superfamily members ^47^, the thiol of the CoA portion protrudes between a V-shape gap, between 2 parallel beta-strands (β18 and β20) of *Ct*NAT10 (**Supplemental Figure 7**).

Although the cryo-EM density for the cytidine-CoA probes clearly show the CoA portion, the linkers and cytidine nucleotide is more poorly resolved (**Figure 4B**). Detailed inspection of this area reveals that the amide linker makes van der Waals interactions with the backbone of Val636 and hydrogen bond interactions with the sidechain -OH of Ser726 and the amide -NH, but no protein interactions are made with the cytidine portion (**Figure 4A**). This observation apparently explains the relatively poor density for the cytosine base, with no density observed at all for of the terminal sugar ring, suggesting that the nucleotide is more flexibly positioned. This greater flexibility of the cytidine nucleotide could suggest that other regions of the RNA substrates are required for appropriately positioning the cytosine base for acetylation.

Inspection of the active site proximal to the cytidine nucleotide reveals a somewhat open pocket with no residues in proximity to make direct contacts suggesting that a structural rearrangement may be required for N4-cytidine acetylation. Consistent with this possibility, the only two residues within ∼13 Å of the cytidine N4 that could play a role in chemistry are His548 and Tyr549, which sit on a loop between helices α18 and α19 of the GNAT domain. Interestingly, this loop sits against several loops within the helicase domain, where a structural change in the helicase domain (due to ATP hydrolysis, RNA substrates binding, or both) could propagate to the loop harboring His548 and Tyr549 (**Figure 4C**). Consistent with the potential catalytic importance of these resides, they are evolutionarily conserved within the eukaryotic NAT10 and TmcA proteins (**Supplemental Figure 2**) and supplied with mutant protein of the corresponding residues in TmcA to alanine were poorly able to rescue ac^4^C formation on total RNA in a *E.coli tmc*AΔ strain ^40^. To directly evaluate the importance of His548 and Tyr549 in catalysis, we characterized the thermal stability and catalytic activity of alanine mutations in both positions and found that while thermal stability was unchanged both mutants showed no detectable catalytic activity, thus supporting their potential roles in catalysis (**Figures 4D-F** and **Supplemental Figures 8**).

### Electropositive patches on the NAT10 surface participate in RNA acetylation

To probe for potential RNA binding surfaces on *Ct*NAT10, we targeted three electropositive patches for mutagenesis for acetyltransferase activity and RNA binding using the h45 rRNA substrate (**Figure 4D, Supplemental Figures 8 and 9**). Surface 1 was located on the front side of the GNAT domain and most proximal to the active site, surface 2 was located on the RNA binding domain of the opposing subunit and located about 20 Å away from the active site and surface 3 was located on the back side of the GNAT domain surface of the opposing subunit and located about 60 Å away from the active site (**Figure 4D**). Both sites 2 and 3 were within distances that could participate in the binding of the extended tRNA and rRNA molecules. Surprisingly, we found that while mutations within all three patches had no effect on protein integrity and thermal stability, (**Figure 4E, Supplemental Figure 8A and 8B**), the mutations significantly compromised acetyltransferase activity, while the more distal patch 3 showing the greatest effect (**Figure 4F and Supplemental Figure 8C**). Interestingly, none of the mutations significantly compromised RNA binding capability (**Supplemental Figure 9**). Together with the fact that the mutations did not affect the binding of substrate acetyl-CoA (**Figure 4E**), these results suggest that the precise mode of RNA binding, rather than the strength of binding is required for acetylation by NAT10.

### Putative catalytic and RNA binding mutants exhibit temperature sensitivity in yeast

To understand whether these biochemical observations extended to living organisms, we developed a functional complementation assay to assess the *in vivo* effects of NAT10 mutations in yeast. This assay is based on a previously developed strain of *Saccharomyces cerevisiae*, where endogenous NAT10 (*Sc*Kre33) has been deleted and rescued with a constitutively expressed inactive mutant (Δkre33+*Sc*H545A), which grows normally under optimal conditions (30 °C) but exhibits decreased fitness at elevated temperatures (37 °C; **Figure 5A**) ^9^. Our hypothesis was that ectopic expression of catalytically active *Sc*Kre33 would rescue this growth defect, while inactive variants would not. Since *Ct*NAT10 and *Sc*Kre33 exhibit high homology, we used our biochemical studies to guide the design of expression vectors encoding wild-type (WT) *Sc*Kre33 as well as several putative acetyltransferase activity-disrupting variants. As expected, we observed that WT-*Sc*Kre33 increases the growth of this strain at 37 °C while an empty vector does not (**Figure 5B**). Transfections with mutant vectors corresponding to putative catalytic mutants of *Ct*NAT10 (*Sc*H545A/*Ct*H548A and *Sc*Y546A/*Ct*Y549A) as well as mutants in basic patches 1 and 3 failed to rescue the temperature sensitivity, while basic patch 2 mutants did show some rescue (**Figure 5B and Supplemental Figure 10**).

To confirm that rescue of growth correlated with rescue of RNA acetylation, we applied nucleotide resolution ac^4^C sequencing to the total RNA extracted from the Δkre33+*Sc*H545A (corresponding to putative catalytic residue *Ct*H548A) and Δkre33+*Sc*R637A (corresponding to a basic patch 1 residue *Ct*R637A) strain ^48^. Assessing 18S rRNA helix 45 (*Sc*-h45), we observed efficient rescue of ac^4^C1773 by the WT plasmid. However, neither mutant had any effect (**Figure 5C and Supplemental Figure 11**). These data specify the importance of H548 and R637 to RNA acetyltransferases across evolution and emphasize the importance of NAT10’s catalytic activity to eukaryotic cell growth.

### Catalytic activity of NAT10 promotes cellular senescence in human cells

Previous studies have demonstrated that NAT10 promotes cellular senescence in colorectal cancer cells ^49^. To evaluate if putative catalytic and RNA binding mutants of NAT10 would exhibit a similar phenotype, we utilized an etoposide induced senescent model in primary lung fibroblast, IMR-90 cells ^50^. NAT10 knockdown IMR90 cells were generated using short hairpin RNAs (shRNAs) targeting endogenous NAT10 (**Figure 6A**). Interestingly, we found that the depletion of NAT10 decreased etoposide-induced senescence and was accompanied by a reduction in senescence markers p16, and secretory senescence-associated phenotype (SASP) markers IL8 and MMP3 (**Figure 6B**). This indicates that attenuation of NAT10 can mitigate senescence in this model system. To further explore this phenomenon, we assessed whether increased expression of NAT10 can exacerbate senescence. We engineered an NAT10 overexpressing cell line (NAT10^WT^-OE) and upon treatment with etoposide observed an increase of senescence by a significant upregulation of mRNA levels of senescence genes, p16, IL8, and MMP3 (**Figure 6C and 6D**). Additionally, we observed a distinct decrease in LaminB1 expression, another hallmark of senescence. Taken together, our findings demonstrate a strong positive correlation between NAT10 expression and senescence regulation.

**Figure 6.**
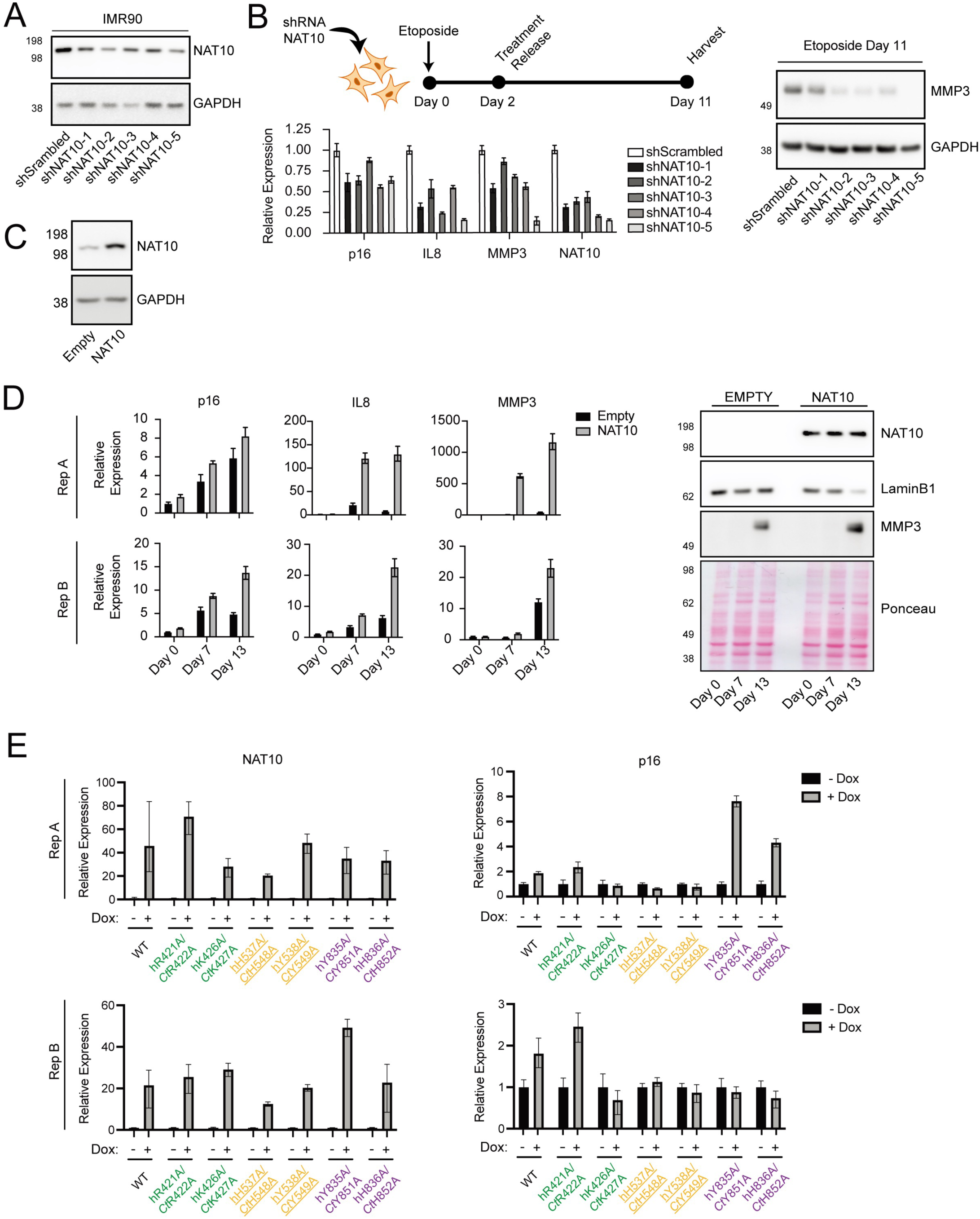
Role of human NAT10 and mutants in cellular senescence in human cells. **(A)** Immunoblot of IMR90 cells depleted of NAT10 via shRNA. GAPDH is used as a loading control. **(B)** Top - Schematic of etoposide-induced senescence. NAT10 depleted IMR90 were treated with 100 µM of etoposide for 48 hours and recovered in untreated media for an additional 9 days. Bottom - RT-PCR of NAT10 and senescence markers during etoposide induced senescence. Values are normalized to shScrambled control. Right - Immunoblot of MMP3 during etoposide induced senescence. GAPDH is used as a loading control. Error bars represent standard deviation of three technical replicates. **(C)** Immunoblot of IMR90 cells overexpressing NAT10 or empty control. GAPDH is used as a loading control. **(D)** IMR90 overexpressing NAT10 or empty control were treated with 100 µM of etoposide for 48 hours and recovered in untreated media and harvested on Day 7 and Day 13. Left - RT-PCR of NAT10 and senescence markers during etoposide induced senescence time course. Values are normalized to Day 0. Right - Immunoblot of Lamin B1 and MMP3 during etoposide induced senescence. Ponceau is used as a loading control. Error bars represent standard deviation of three technical replicates. **(E)** RT-PCR of p16 in IMR90 cells expressing either WT or mutant NAT10 under tetracycline induction. Cells were treated with and without doxycycline and undergo etoposide treatment for 48 hours and recovered in undertreated media for an additional 8 days. Values are normalized to non-dox treated cells. Residues are color-coded according to the regions in Figure 4D. Putative catalytic residues (hH537A/*Ct*H548A and hY538A/*Ct*Y549A) are underlined. Error bars represent standard deviation of three technical replicates.

Having substantiated NAT10’s ability to promote senescence in this model, we next aimed to evaluate the involvement of NAT10’s acetyltransferase activity in this process. To ask this question, we generated IMR90 cell lines with tetracycline inducible expression of either wild-type hNAT10 or point alanine mutants corresponding to conserved residues determined in our structural and biochemical studies to be essential for NAT10 activity. Consistent with the above results, expression of WT-NAT10 showed increased expression of the senescence marker p16. In contrast, the hNAT10 mutants hK426A, hH537A, and hY538A (corresponding to *Ct*K427A-patch 3 mutant, *Ct*H548A-active site mutant, and *Ct*Y549A-active site mutant) exhibited a markedly decreased p16 expression, while the rest of the mutants recapitulated the effect of WT-NAT10 (**Figure 6E**). Interestingly, one patch 3 mutant (hR421A/*Ct*R422A) and patch 2 mutants (hY835A/*Ct*Y851A and hH836A/*Ct*H852A) in both the human cell-based senescence assays and the yeast cell-based fitness assay scored as WT or near WT, but were either inactive or impaired in *Ct*NAT10 biochemical assays, suggesting some differences between the *in vitro* and cellular systems. Nonetheless, overall, these studies implicate NAT10’s acetyltransferase domain and catalytic activity as a functional driver of cellular senescence.

## DISCUSSION

In all eukaryotic systems that have been evaluated to date, RNA acetyltransferases are essential for life. However, the contribution of individual domains to NAT10-dependent biology has been difficult to understand. Here we report the first cryo-EM structure of a stand-alone eukaryotic RNA acetyltransferase (*Ct*NAT10) bound to CoA-cytidine bisubstrate probes in the presence and absence of ADP. The structures reveal a symmetrical, heart-shaped dimer that is distinct from the L-shaped monomer of the orthologous bacterial TmcA enzyme ^40^. Our analysis reveals a unique active site, with the cytidine base of the bisubstrate probe surrounded by conserved GNAT, helicase, DUF and RNA binding domains (from the opposing subunit of the dimer). Structure-based mutagenesis validated the importance of two conserved active site residues (*Ct*H548 and *Ct*Y549) as well as three basic patches on the dimer surface in RNA binding and cognate nucleic acid acetylation. These interactions appear to be important to RNA acetyltransferase activity across evolution, and were used in yeast and human cell lines to identify a role for NAT10 catalysis thermoadaptation and cellular senescence, respectively.

The GNAT domain of *Ct*NAT10, which anchors the cytidine-linked CoA probe, contains the same overall structure of the related GNAT folds as lysine-targeting histone acetyltransferases (HATs) and N-terminal acetyltransferases (NATs). However, a structural superposition to the three folds reveals that the substrate binding tunnel of *Ct*NAT10 is wider than that of HATs and NATs, consistent with the need to acetylate larger and more highly charged RNA substrates (**Figure 7**).

**Figure 7.**
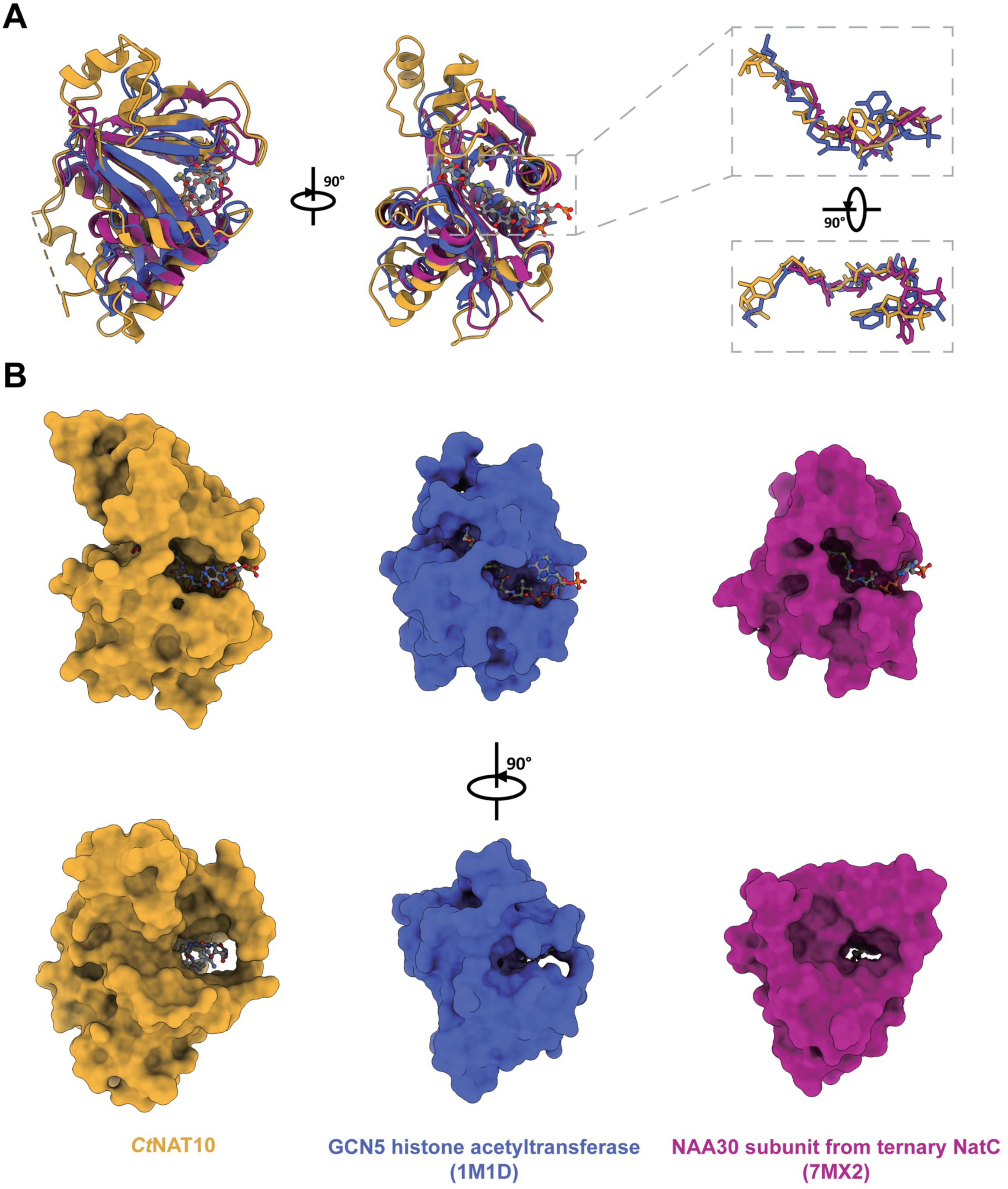
Architecture of the substrate binding tunnel among GNAT family members. **(A)** Superposition of GNAT domain of *Ct*NAT10 with representative Histone acetyltransferase (GCN5) and N-terminal acetyltransferase (NAA30) proteins. *Ct*NAT10 (bound to cytidine-amide-CoA) is colored in yellow, *Tetrahymena thermophilus* GCN5 (bound to Lys-CoA, PDB: 1MID ^44^) is colored in blue, and human NAA30 (bound to peptide-CoA, PDB: 7MX2 ^73^) is colored in magenta. In the structure alignment, only cytidine-amide-CoA is shown as ball-and-stick representation to better demonstrate bisubstrate CoA binding groove. The right panel shows the conformational comparison among 3 bisubstrate CoA analogs when the Cα of GNAT domains are aligned. **(B)** GNAT domain surfaces of *Ct*NAT10, GCN5 and NAA30 are shown in both front and side views corresponding to the color code in A, with their in-complexed bisubstrate CoA probes shown as ball-and-stick representation, respectively.

In an attempt to identify potential RNA binding surfaces on *Ct*NAT10 for rRNA and tRNA substrates, we targeted selected basic patches (called 1, 2 and 3, see **Figure 4D**) on *Ct*NAT10 for mutagenesis, followed by an assessment of catalytic and RNA binding activities. Surprisingly, we found that while mutations within two of the three patches (1 and 3) significantly decreased catalytic activity, these same mutations did not impair RNA binding activity. Of note, the NAT10 surface contains other basic patches that could participate in non-specific RNA binding, that could compensate for a loss of RNA binding affinity at specific binding sites, that may be required for correctly positioning the cytidine base within the NAT10 binding site for acetylation. We therefore hypothesize that the precise binding mode, rather than overall binding strength, is critical for enzymatic activity. NAT10 enzymes have been reported to work together with Tan1/Thumpd1 cofactors for tRNA acetylation ^9,21^, and several snoRNAs such as snR4, snR45/SNORD13 for rRNA acetylation in yeast and human, respectively ^22-25^. These additional macromolecules may also play a role in RNA recognition, and potentially also catalysis, by NAT10 enzymes.

Our ability to identify NAT10 mutants that were impaired in catalytic activity *in vitro* allowed us to characterize the effect of these mutants in cellular models of normal yeast growth under heat stress and markers of cellular senescence in human cells. We found that the alanine mutation of two residues in the active site that are important for catalysis *in vitro* (*Ct*H548 and *Ct*Y549) in the NAT10, exhibited a temperature sensitivity in yeast and inhibited senescence in human cells, in both cases phenocopying NAT10 deletion. Given the additional non-catalytic role of NAT10 in ribosome biogenesis, these results point to the particular importance of the catalytic cytidine N4 acetylation activity of NAT10 to the temperature sensitivity in yeast and senescence phenotype in human cells. While the specific pathways that link the cytidine N4 acetylation activity of NAT10 to these phenotypes is beyond the scope of this study, these findings do highlight the potential utility of targeting the catalytic function of NAT10 in senescence and ageing-associated diseases such as HGPS.

## Lead Contact

Further information and requests for resources and reagents should be directed to and will be fulfilled by lead contact, Ronen Marmorstein.

## Materials Availability

Plasmids and strains generated in this study are available upon request from the corresponding authors.

## Data And Code Availability

The cryo-EM maps and atomic coordinates for *E.coli* produced recombinant *Ct*NAT10/cytidine-acetone-CoA, *E.coli* produced recombinant *Ct*NAT10/cytidine-amide-CoA/ADP and *Sf9* produced recombinant *Ct*NAT10/cytidine-amide-CoA/ADP have been deposited to the Electron Microscopy Data Bank (EMDB) (accession codes: 43996, 44038 and 44042, respectively) and the Protein Data Bank (PDB) (accession codes: 9AYM, 9B0E and 9B0I, respectively). Primary data are available upon request from the corresponding authors.

Any additional information required to reanalyze the data reported in this paper is available from the lead contact upon request.

## Inclusion And Diversity

We support inclusive, diverse, and equitable conduct of research.

## Acknowledgement

We thank Dr. Darrah Johnson-McDaniel, Dr. Stefan Steimle and Dr. Sudheer Molugu at the Perelman School of Medicine, University of Pennsylvania for helping with cryo-EM data collection, Dr. Ying Xiong (NCI) for assisting with synthetic characterization data, and Sunny Sharma (Rutgers University) for providing the pPK468 plasmid. Funding for this project was provided by NIH grant R35GM11890 to RM, ZIABC011488 to JLM and P01AG31862 to RM and SB. Molecular graphics and analyses are mostly performed with UCSF Chimera, developed by the Resource for Biocomputing, Visualization, and Informatics at the University of California, San Francisco, with support from NIH P41-GM103311.

## Author Contributions

MZ, STG, DB, KT, SB, JLM and RM conceptualized and designed the project. MZ, STG DB, KT, XW and BY carried out experiments. SB, JLM and RM supervised the project. MZ wrote the first draft of the manuscript. All authors edited and approved the final version of the manuscript.

## Experimental Model And Subject Details

For biochemical experiments and cryo-EM structural analyses, we employed *Spodoptera frugiperda* (*Sf9*) cells grown in SFM II media for the recombinant expression of *Ct*NAT10 protein. We also used homemade BL21 competent *E.coli* supplemented with a plasmid expressing HSP70 (bGroESL) to prepare recombinant *Ct*NAT10 for cryo-EM structural studies.

## Key Resource Tables

**Table S1.**
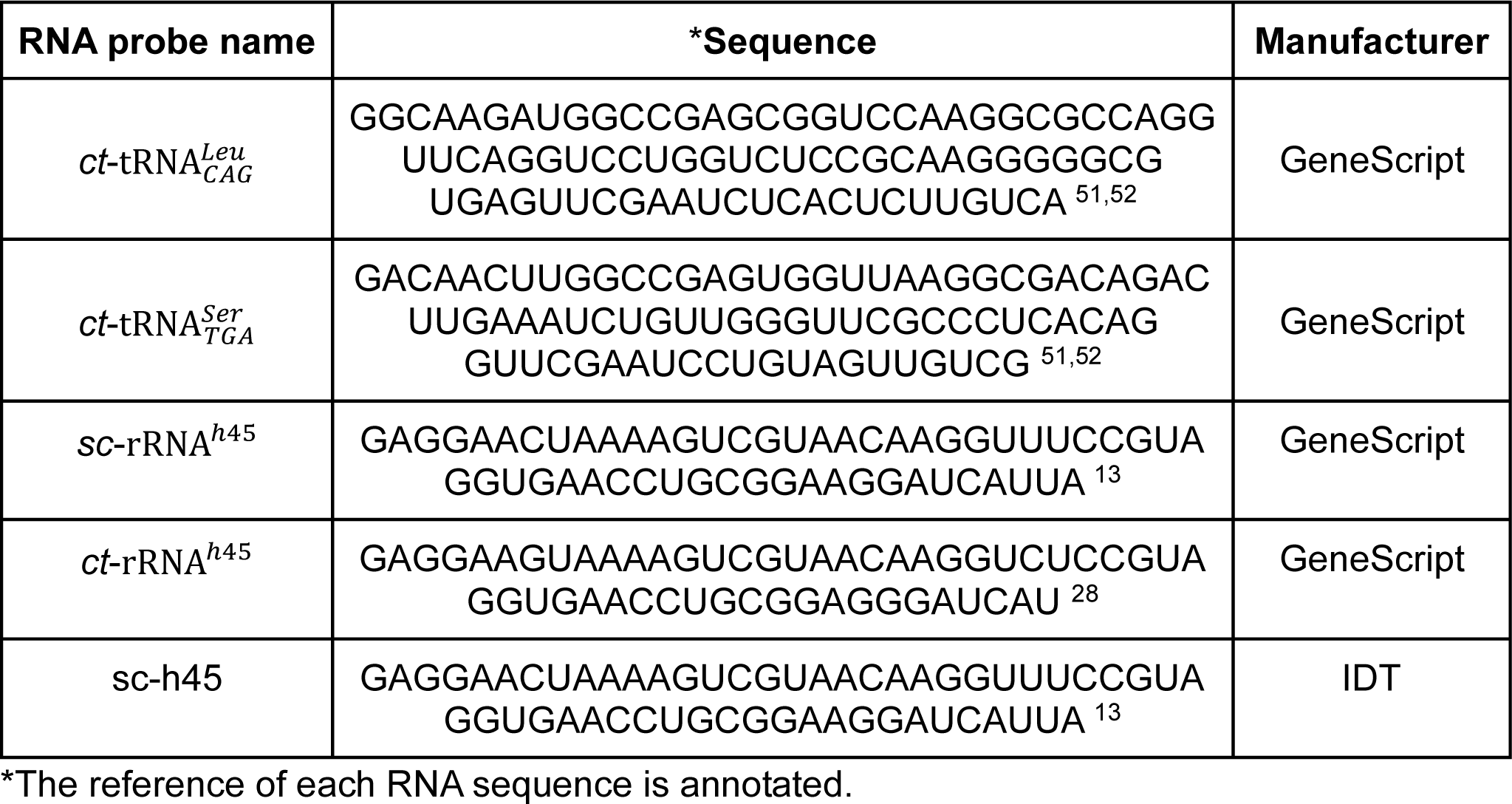
synthesized RNA for EMSA and *In vitro* RNA acetylation assay.

**Table S2.**
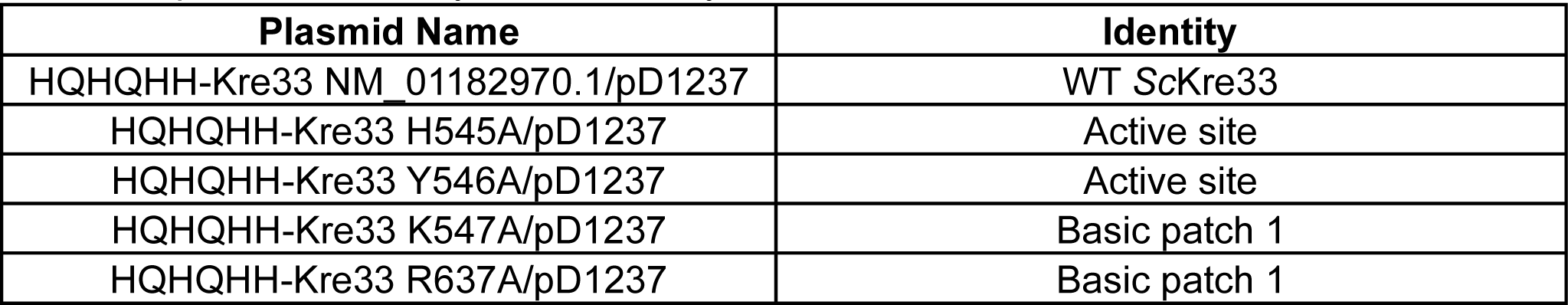
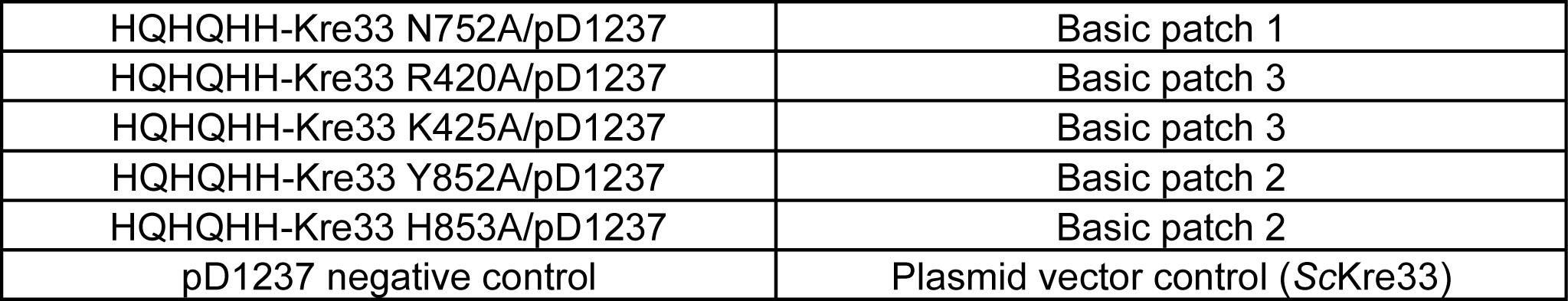
plasmid used in yeast cell assays.

**Table S3.**
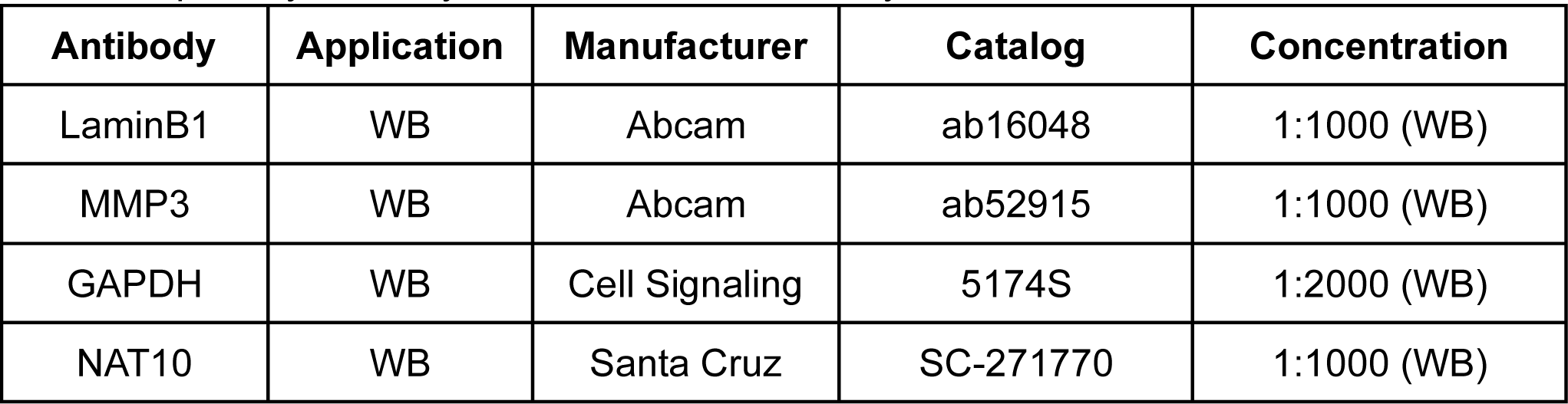
primary antibody used in human cell assays.

**Table S4.**
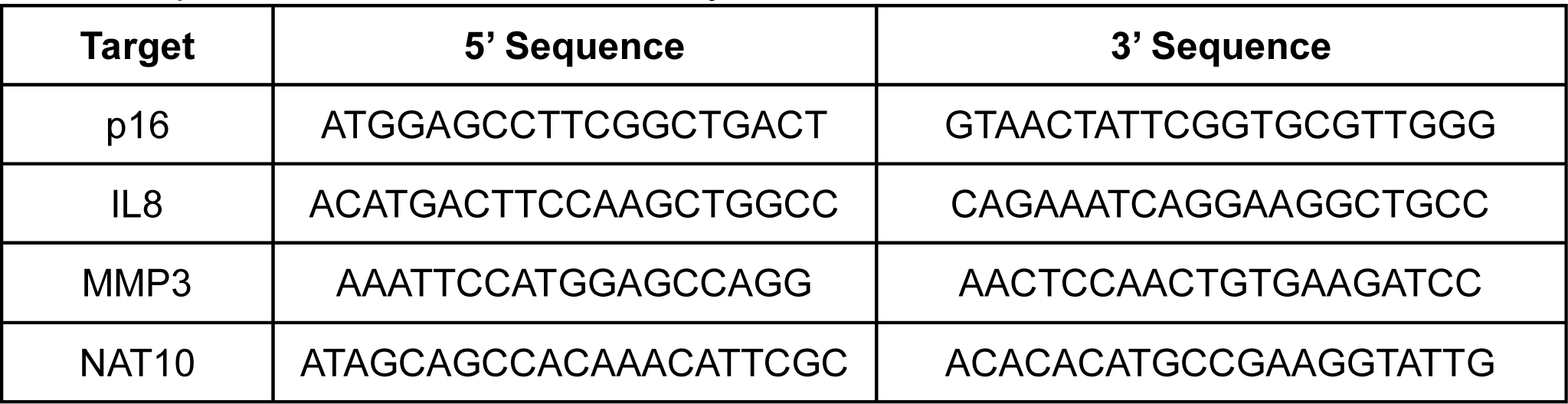
primer used in human cell assays.

## METHODS

### *Ct*NAT10 expression and purification

A gene encoding N-terminal 6xHis tagged WT-*Ct*NAT10 was synthesized, and codon optimized for overexpression in both *Sf9* and bacteria (BioBasic). Recombinant *Ct*NAT10 protein was expressed in both *Sf9* cells (ThermoFisher, cat #12659017) and homemade bGroESL cells (See **Experimental Model and Subject Details**). For *Ct*NAT10 expression in *Sf9*, the synthesized gene was subcloned into a pFastBacHT A vector (WT-*Ct*NAT10-pFastBacHT A plasmid). High density (2 × 10^6^ cells * ml ^−1^) suspension cultures of *Sf9* cells were infected at a multiplicity of infection of ∼1, and incubated for 48 h in Fernbach Shake flasks at 27 °C. All subsequent purification steps were carried out at 4 °C. Cell pellets were harvested and per liter of cell culture was resuspended in 50 ml lysis buffer containing 25 mM Hepes-NaOH pH 7.5, 500 mM NaCl, 10% (v/v) glycerol, 10 mM β-mercaptoethanol (β-ME), and an EDTA-free protease inhibitor tablet (Roche). After mild sonication (2-minute sonication time per liter of cell culture, 5 seconds “on”/30 seconds “off” at 40% amplitude, with sonication cup fully buried in ice), 2500 U benzonase nuclease (MilliporeSigma), 200 U RNAse cocktail enzyme mix (Invitrogen) and 0.25 mmol MgCl_2_ were added per liter of cell culture. To ensure that RNAse and benzonase nuclease could sufficiently remove endogenous nucleic acids, the lysate was incubated on a stir plate for 30 minutes. The cell lysate was then clarified with centrifugation and the supernatant was collected and loaded onto a gravity column manually loaded with TALON Metal Affinity Resin (Takara), washed 5 times with 10 column volumes (CV) of wash buffer containing 25 mM Hepes-NaOH pH = 7.5, 500 mM NaCl, 15 mM imidazole, 1 mM tris-(2-carboxyethyl)phosphine (TCEP), and eluted with elution buffer containing 25 mM Hepes-NaOH pH = 7.5, 500 mM NaCl, 200 mM imidazole, 1 mM TCEP. The eluent was concentrated using a 100 kDa MWCO centrifugal filter (Amicon Ultra, MilliporeSigma) and loaded onto a Superdex 200 Increase 10/300 GL gel filtration column (GE Healthcare) in sizing buffer containing 25 mM Hepes-NaOH pH = 7.5, 300 mM NaCl, and 1 mM TCEP. Only peak fractions with UV_260/280_ < 0.8 were collected and concentrated to ∼1 mg/ml determined by Nanodrop 2000 (Thermo Fisher Scientific), and aliquoted into 20 μl aliquots and flash-frozen in LN_2_ for storage at −80 °C until use.

For *Ct*NAT10 *E. coli* overexpression, the synthesized gene was subcloned into a pET30a(+) expression vector. Transformed cells were cultured at 37 °C until the optical density reached an 600nm (OD_600_) of ∼0.8, induced with 0.5 mM Isopropyl β-D-1-thiogalactopyranoside (IPTG) at 17°C, and grown overnight (16-20 h). The cell pellet was harvested and resuspended in lysis buffer containing 25 mM Hepes-NaOH pH = 7.5, 500 mM NaCl, 10% (v/v) glycerol, 10 mM β-mercaptoethanol (β-ME), and an EDTA-free protease inhibitor tablet (Roche). The protein was purified to homogeneity using a combination of TALON Metal Affinity Resin (Takara) and gel filtration chromatography, essentially as described above for *Sf9*-expressed WT-*Ct*NAT10.

The WT-*Ct*NAT10-pFastBacHT A plasmid was used to prepare all *Ct*NAT10 mutants using the QuikChange protocol (Stratagene) ^53^. All mutant proteins were overexpressed and purified to homogeneity following the same procedure described above for *Sf9*-expressed WT-*Ct*NAT10.

### Synthesis of cytidine-CoA structural probes

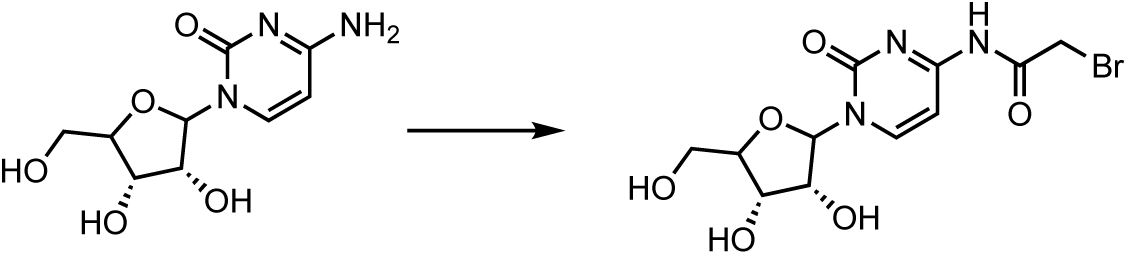

#### N4-bromoacetylcytidine

To a solution of 2-bromoacetic acid (187 mg, 0.71 mmol) in DMF (3 ml) was added diisopropylcarbodiimide (119 μl, 0.71 mmol). After stirring for 5 min at 23 °C, cytidine (187 mg, 0.71 mmol) was added, and the reaction mixture was stirred for another 45 min. DMF was removed in vacuo to provide the crude product as a light pink residue which was purified by silica flash chromatography (8% MeOH in DCM) to yield N4-bromoacetylcytidine as a white solid (107 mg, 38% yield). ^1^H NMR (500 MHz, DMSO) δ 11.23 (s, 1H), 8.48 (d, J = 7.5 Hz, ^1^H), 7.12 (d, J = 7.4 Hz, 1H), 5.78 (d, J = 3.0 Hz, 1H), 4.14 (s, 2H), 4.02 – 3.94 (m, 2H), 3.90 (dt, J = 6.0, 2.9 Hz, 1H), 3.74 (dd, J = 12.2, 2.9 Hz, 1H), 3.60 (dd, J = 12.3, 3.0 Hz, 1H), 3.17 (s, 4H). LRMS (ESI/Single Quad) calcd for C_11_H_15_BrN_3_O_6_ ^+^ [M+H]^+^ 364.0, found 364.1.

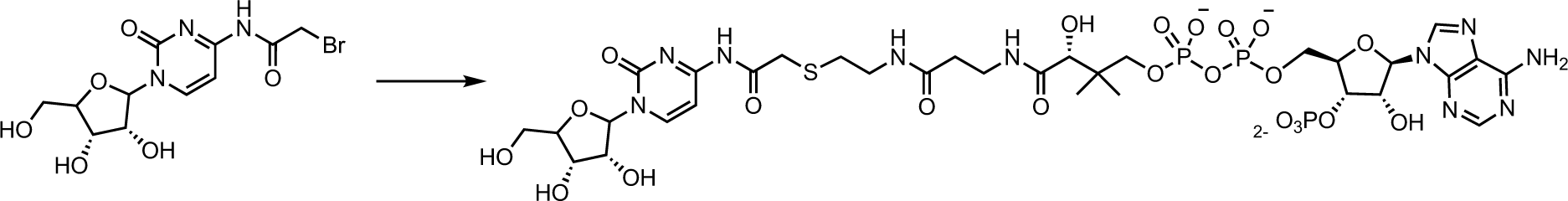

#### Cytidine-amide-CoA

To a solution of N4-bromoacetylcytidine (25 mg, 68 μmol) in DMSO (85 μl) was added coenzyme A (26 mg, 34 μmol) in phosphate buffer (100 mM, 85 μl, pH 7.0). The reaction mixture was stirred at 23 °C for 2.5 h, at which time CoA had been consumed as monitored by HPLC. The crude reaction mixture was diluted with aqueous triethylammonium acetate (TEAA, 0.5 ml, 20 mM, pH 7), and then purified by C18 flash chromatography [Solvent A: TEAA (20 mM, pH 7); Solvent B: acetonitrile. Gradient: 0% B for 2 column volumes (CV), 0% to 15% B over 12 CV, 15% B for 10 CV, 15% to 30% B over 10 CV, 30% to 50% B over 1 CV, and 50% B over 5 CV] to yield the bisubstrate conjugate as a white solid (5.5 mg, 15% yield). ^1^H NMR (500 MHz, D_2_O) δ 8.53 (s, 1H), 8.31 (d, J = 7.6 Hz, 1H), 8.23 (s, 1H), 7.21 (d, J = 7.5 Hz, 1H), 6.13 (d, J = 6.6 Hz, 1H), 5.88 (d, J = 2.5 Hz, 1H), 4.63 – 4.50 (m, 1H), 4.34 – 4.28 (m, 1H), 4.23 (d, J = 3.8 Hz, 2H), 4.19 (d, J = 3.0 Hz, 2H), 4.03 (s, 1H), 4.01 (dd, J = 12.8, 1.9 Hz, 1H), 3.89 – 3.80 (m, 2H), 3.55 (dd, J = 9.7, 4.7 Hz, 1H), 3.51 – 3.43 (m, 3H), 3.38 (t, J = 6.5 Hz, 2H), 2.76 (t, J = 6.6 Hz, 2H), 2.45 (t, J = 6.5 Hz, 2H), 1.92 (s, 6H), 0.89 (s, 3H), 0.76 (s, 3H). 31P NMR (202 MHz, D_2_O) δ 1.83, -10.90 (d, J = 20.8 Hz), -11.48 (d, J = 20.9 Hz). LRMS (ESI/Single Quad) calcd for C_32_H_50_N_10_O_22_P_3_S^+^ [M+H]^+^ 1051.2, found 1050.6. While the resulting compound is N4-acetylcytidine-amide-CoA, we refer to it in the text as Cytidine-amide-CoA for simplicity.

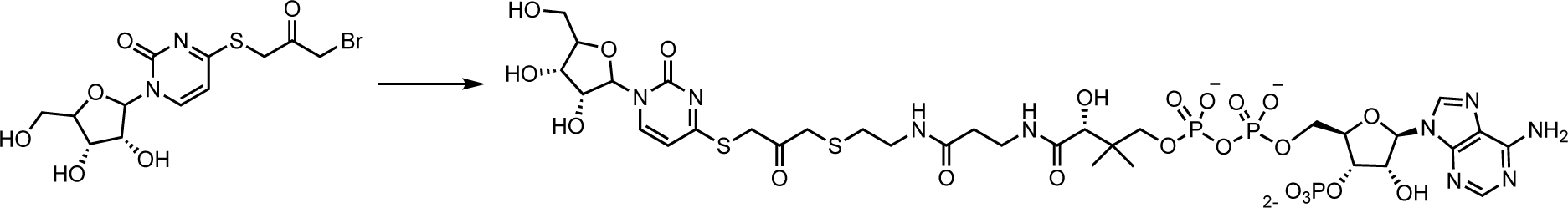

#### Cytidine-acetone-CoA

To a solution of bromoacetonyl-thiouridine (25 mg, 63 μmol) in DMSO (85 μl) was added coenzyme A (49 mg, 63 μmol) in phosphate buffer (100 mM, 3 ml, pH 7.0). The reaction mixture was stirred at 23 °C for 15 min. The crude reaction was then diluted with 0.1 % trifluoroacetic acid (TFA) and purified by reverse phase HPLC (0.1% TFA/CH_3_CN) to yield the acetone-linked CoA probe. ^1^H NMR (500 MHz, D_2_O) δ 8.57 (s, 1H), 8.34 (s, 1H), 8.22 (s, 1H), 6.68 (s, 1H), 6.16 – 6.09 (m, 1H), 5.75 (d, *J* = 2.4 Hz, 1H), 4.82 – 4.72 (m, 3H), 4.52 – 4.48 (m, 1H), 4.16 (dddd, *J* = 17.0, 11.7, 4.8, 2.9 Hz, 3H), 4.12 – 3.99 (m, 2H), 3.93 (s, 1H), 3.88 (dd, *J* = 12.9, 2.2 Hz, 1H), 3.80 – 3.65 (m, 2H), 3.63 (s, 2H), 3.56 – 3.47 (m, 1H), 3.38 (t, *J* = 6.5 Hz, 2H), 3.24 (t, *J* = 6.6 Hz, 2H), 2.59 (t, *J* = 6.6 Hz, 2H), 2.37 (t, *J* = 6.5 Hz, 2H), 0.84 (s, 2H), 0.70 (s, 3H). ^13^C NMR (126 MHz, D_2_O) δ 177.31, 174.70, 173.92, 149.85, 148.47, 144.63, 142.48, 141.75, 118.52, 91.69, 87.42, 83.82, 83.49, 74.40, 74.18, 74.14, 74.08, 73.92, 71.90, 68.34, 65.08, 59.88, 39.15, 38.35, 38.28, 38.06, 35.41, 35.32, 31.34, 30.82, 20.84, 18.16. LRMS (ESI/Single Quad) calcd for C_33_H_51_N_9_O P S ^+^ [M+H]^+^ 1082.2, found 1082.8. While the resulting compound is thiouridine-acetonyl-CoA, we refer to it in the text as Cytidine-acetone-CoA for simplicity.

### Differential scanning fluorimetry assay

To investigate the thermal stability of *Ct*NAT10 alone or in the presence of different cofactors, individual 20 μl reactions were prepared containing 1μM of *ct*NAT10 and a 1:1000 dilution of a 5000× concentrated SYPRO orange dye (Thermo Fisher Scientific) in DSF buffer (37.5 mM Hepes-NaOH pH = 7.5, 150 mM NaCl, 6 mM KCl, 5 mM MgCl_2_, 0.5 mM DTT, 0.5 mM TCEP). Three replicate samples were prepared for each combination of protein and additives with the indicated concentration in the figure legends, and distributed to separate wells in a 384-well assay plate specifically designed for real-time qPCR measurements. Thermal melting curves were obtained by heating the samples from 20 to 95 °C using a ramping rate of 2%, with fluorescence detection at 580 nm, monitored using a 7900HT Fast Real Time PCR System (Applied Biosystems). The fluorescence was normalized based on the peak value, and Tm was defined as the temperature at the inflection point of the SYPRO fluorescence curve. To assess the variability of the results, we calculated the standard deviation of each condition and plotted error bars. Plot generation and P-values calculation was performed using GraphPad Prism version 9 for Windows, GraphPad Software, Boston, Massachusetts USA, www.graphpad.com.

### Electrophoretic mobility shift assays (EMSA)

For testing the purified *Ct*NAT10 wild-type binding with synthesized tRNA and rRNA substrates, all the RNA probes were synthesized, purified by RNase free HPLC, and dissolved in DNase/RNase-free water at 1 mM by Genescript (for sequence detail see **Key Resources Table S1**). To conduct binding assays, samples containing 0.5 μM of the indicated RNA substrate with various concentrations (indicated in the figure legends) of purified *Ct*NAT10 proteins were incubated overnight (15 h) at 4°C in a buffer containing 50 mM Hepes-KOH pH = 7.6, 90 mM NaCl, 12 mM KCl, 10 mM MgCl_2_, 1 mM DTT, 0.3 mM TCEP, 1.0 U/μl SUPERase-In RNAse inhibitor (Invitrogen), 0.5 mM ATP, and 0.1 mM acetyl-CoA. The next day, samples were incubated for 1.5 h at 45 °C, and the entire 10 μl RNA–protein mixture was then loaded onto DNA Retardation Gels (6%) (Invitrogen) with 2.5 μl Novex™ Hi-Density TBE Sample Buffer (5X) (Invitrogen). The gel was run in 0.5x TBE buffer at 120 V for 42 min and stained with 1x SYBR™ Gold Nucleic Acid Gel Stain (Invitrogen) diluted with 0.5x TBE buffer for 20 min. Imaging was performed using a ChemiDoc Imaging System (BIO-RAD), and quantification was carried out using Fiji ImageJ ^54^ to measure the intensity of the free RNA band. For protein band visualization, the same gels were subjected to three Milli-Q water washes to remove excess SYBR Gold, and subsequently stained with Coomassie Blue for 20 minutes, followed by de-staining overnight with agitation in Milli-Q water.

For evaluating *Ct*NAT10 alanine mutant binding with h45 RNA substrate, A *Saccharomyces cerevisiae* (sc) 18 S rRNA helix 45 (sc-h45) probes (sequence detail see **Key Resources Table S1**) was synthesized and purified by RNase-free HPLC by IDT Corporation. Lyophilized sc-h45 powder was dissolved in DNase/RNase-free water at 100 μM and the final concentration of sc-h45 in each sample was 0.25 μM. The binding assays and RNA band intensity measurement were carried out essentially as described above. Dissociation constant (K_d_) calculation and binding curve fitting was performed using GraphPad Prism version 9 for Windows, GraphPad Software, Boston, Massachusetts, USA, www.graphpad.com. Specific binding with Hill slope equation (Y=B_max_ * X^h/(K_d_^h + X^h) was use for curve fitting, where Y is the fraction complex, X is the concentration of *Ct*NAT10 (in micromolar), B_max_ is the maximum binding and h is the Hill slope. Three technical replicates were performed for all the mutant EMSAs.

### *In vitro* RNA acetylation assay

Assays were performed largely following the methods described previously ^13^, with some modifications. Briefly, reaction mixtures (100 µl) containing 50 mM Hepes-KOH pH = 7.6, 60 mM NaCl, 12 mM KCl, 10 mM MgCl_2_, 1 mM DTT, 1.0 U/μl SUPERase-In RNAse inhibitor (Invitrogen), 0.5 mM ATP, 0.1 mM [1-^14^C] acetyl-CoA (PerkinElmer, 60 mCi/mmol), 5 µM sc-h45 RNA, and 3 µM recombinant WT or mutant *Ct*NAT10 were incubated at 45 °C for the times indicated in the figure legends. After reaction, the substrate RNAs were extracted by phenol: chloroform: isoamyl alcohol (25:24:1, v/v) (Invitrogen), precipitated by ethanol, and analyzed by 10% TBE-Urea Gel (Invitrogen). The substrate RNAs were stained with 1x SYBR™ Gold Nucleic Acid Gel Stain (Invitrogen) diluted with 1x TBE buffer for 10 min. Gels were visualized using a Typhoon™ laser-scanner platform (Cytiva). Subsequently, the gels were dried *in vacuo*, and the radioactivity of the acetylated RNA fragments were exposed to a phosphor screen overnight and visualized using a Typhoon™ laser-scanner platform. Band intensities were quantified using Fiji ImageJ ^54^, and the relative activity of *Ct*NAT10 mutants were normalized to the WT-*Ct*NAT10 activity. Three technical replicates were performed for all the activity assays.

### Mass photometry

Mass photometry sample slides were manually assembled by placing a 6-wells silicone cassette onto the sample carrier cover glass. Samples and standard proteins were diluted with dilution buffer (300 mM NaCl, 25 mM Hepes-NaOH pH = 7.5, 1 mM TCEP) at room temperature. The dilution buffer was freshly filtered by a 0.22 µm syringe filter before the experiment. For calibration, 2ul of homemade BAM-TG protein standard mix (300 nM thyroglobulin, 1 μM beta-amylase in PBS with 0.5% glycerol) was added into the droplet of 18 μl dilution buffer and mixed thoroughly before measurement. Before testing each protein sample, 19 μl dilution buffer was loaded into sample well for finding focus and quality metrics checking, then 1 μl of 200 nM protein stock was added into the sample droplet of 19 μl dilution buffer and mixed thoroughly before measurement. For data analysis, a calibration curve was created by manually putting in the molecular weight of the Gaussian fit for each peak of corresponding standard protein (thyroglobulin: 670 kDa, beta-amylase: 224 kDa and 112 kDa) to make sure the R^2^ value for the calibration curve >0.999. Movies for each protein sample were acquired in 60 seconds (3,000 frames in total) with all the other settings at the default values. All data collection was performed using a commercial TwoMP mass photometer with the software AcquireMP, version 2022 R1 (Refeyn Ltd.). Data analysis was performed by DiscoverMP (Refeyn Ltd.).

### Cryo-EM sample preparation and data collection

For all *E.coli* and *Sf9* produced *Ct*NAT10 with cytidine-CoA conjugate compound samples, freshly purified protein was first diluted with sizing buffer (25 mM Hepes-NaOH pH = 7.5, 300 mM NaCl, and 1 mM TCEP) to 0.8 mg/ml. For *E.coli* produced *Ct*NAT10 with the cytidine-acetone-CoA conjugate sample, the protein was incubated with 2 mM of the cytidine-acetone-CoA and 5 mM TCEP pH = 7.5 on ice for 1h; and for both the *E.coli* and *Sf9* produced *Ct*NAT10 with the cytidine-amide-CoA conjugate sample, protein was incubated with 2.5 mM ATP, 5 mM MgCl_2_, 2 mM of the cytidine-amide-CoA conjugate and 5 mM TCEP pH = 7.5 on ice for 1h. For the preparation of grids, 3 μl of protein-cofactor complex sample was applied to glow-discharged (PELCO easiGlow™) Quantifoil R1.2/1.3 300 mesh copper grids, and a FEI Vitrobot Mark IV was used for blotting (Time=2.5s, force=3) under 100% humidity at 7°C, then grids were plunged frozen in liquid ethane. Grid screening was performed with a FEI Titan Krios transmission electron microscope equipped with a K3 Summit direct detector (Gatan) at a magnification of 105,000x, and the most promising grids were used for data collection under the same magnification at super-resolution mode, with defocus values set from -1.0 to -3.0 μm. The details of cryo-EM data collection parameters are described in **Table 1**.

### Cryo-EM data processing and 3D model reconstruction

All three cryo-EM datasets were processed with similar workflows, described as follow. After importin the data into Relion 3.1.3 ^55^, beam-induced motions were corrected for the original image stacks using MotionCor2 ^56^, while the pixel was binned 2-fold. After motion correction, all micrographs were imported into Cryosparc2 ^57^, where they were processed further. CTFFIND4 ^58^ was used to determine the defocus value of each micrograph and micrographs with contrast transfer function (CTF) information poorer than 5 Å were discarded. For the particle picking step, blob picker was utilized to pick ∼50,000 particles from ∼200 micrographs, with a “blob” diameter of 100-200 Å. After detailed particle inspection, selected particles were used for reference-free 2D classifications, and “good” classes were chosen as a “template” for template picking on entire datasets. About 3,000,000 particles were picked for each dataset and went through several rounds of 2D classification to remove the particles that did not appear to be protein particles. Only the best 2D classes with different orientations were selected and 80,000 particles were randomly selected by algorithm to perform *Ab-initio* modeling.

For 3D models reconstruction, 4 initial models were created using *Ab-initio* modeling. All the “good” particles that were selected by 2D classifications went through two rounds of heterogeneous refinement using all 4 initial models as input volume, to further remove junk particles. Maps with overall resolutions of 3.37 Å, 3.56 Å and 3.37 Å were generated for *E.coli* produced recombinant *Ct*NAT10/cytidine-acetone-CoA, *E.coli* produced recombinant *Ct*NAT10/cytidine-amide-CoA/ADP and *Sf9* produced recombinant *Ct*NAT10/cytidine-amide-CoA/ADP datasets, respectively, with no symmetry imposed during initial homogeneous refinement. The map resolutions were further improved to 3.35 Å, 3.48 Å and 3.23 Å after non-uniform refinement ^59^, per-particle global CTF refinement and local CTF refinement. The final maps with overall resolutions of 3.29 Å, 3.19 Å and 3.02 Å was generated with C_2_ symmetry applied.

### Cryo-EM model building and refinement

For the *E.coli* produced recombinant *Ct*NAT10/cytidine-acetone-CoA dataset, the density map was reconstituted from 160,385 particles chosen from 5,500 raw electron micrographs, and the *Ec*TmcA crystal structure (PDB: 2ZPA) was utilized as the initial model for building the *Ct*NAT10 structure. The residues in the *Ec*TmcA structure model were first mutated to corresponding residues of *Ct*NAT10 according to sequence alignment by CHAINSAW ^60,61^ (CCP4 supported program ^62^), then fit into the map as a rigid body utilizing UCSF Chimera ^63^. The atomic coordinates was manually built using COOT ^64^ according to the cryo-EM map, while guided by the secondary structure prediction from XtalPred ^65^. For cytidine-acetone-CoA ligand model building, the initial chemical structure was manually depicted by “Ligand Builder” in COOT ^64^ and roughly placed into the map based on the density. The geometry restraint information was generated by phenix.elbow ^66^, and the information was provided to phenix.real_space_refine program during protein structure auto refinement. The complete model was auto refined using real-space refinement in PHENIX then manually adjusted with COOT ^64^ according to the model evaluation by MolProbity ^67^ for several rounds, until no significant outliers were reported in the phenix.real_space_refine program. The structure is well ordered with all four functional domains well resolved, except for residues 1-5, 63-93, 437-466, 498-510, 667-704, 929-1073, which were not visible in cryo-EM map.

For the *E.coli* produced recombinant *Ct*NAT10/cytidine-amide-CoA/ADP and *Sf9* produced recombinant *Ct*NAT10/cytidine-amide-CoA/ADP datasets, the structure were build using the *E.coli* produced *Ct*NAT10/cytidine-acetone-CoA structure as the starting model, and refined as described above. Both structures are well ordered. For *E.coli* produced recombinant *Ct*NAT10/cytidine-amide-CoA/ADP structure, residues 1-5, 63-93, 437-466, 498-510, 667-695 and 929-1073, were not visible in the cryo-EM map. For *Sf9* produced recombinant *Ct*NAT10/cytidine-amide-CoA/ADP structure, residues 1-4, 64-93, 437-466, 498-510, 667-704 and 930-1073, were not visible in the cryo-EM map.

All representations of cryo-EM density and structural models were prepared with UCSF ChimeraX ^68^ and PyMol ^69^. The dimer interface area between two helicases domain and RNA binding domain were calculated using ‘Protein interfaces, surfaces and assemblies’ service PISA at the European Bioinformatics Institute. (http://www.ebi.ac.uk/pdbe/prot_int/pistart.html) ^70^.

### Yeast cell transformation

Yeast cell transformation with each plasmid construct was performed as previously described ^71^. Briefly, a single colony of Δkre33 parent cells was inoculated in 5 ml of YPD medium and incubated on a rotary shaker at 200 rpm at 30 °C for 16-20 h. To a flask containing 50 ml YPD medium, 2.5 x 10^8^ cells (a suspension containing 1 x 10^7^ cells ml^-1^ will give an OD_600_ of 1) were added to obtain a titer of 5 x 10^6^ cells ml^-1^, and the flask was incubated at 30 °C, 200 rpm until the titer reached at least 2 x 10^7^ cells ml^-1^. Cells were harvested by centrifuging for 5 min at 3000 g. The pellet was washed 2 times by resuspending in 25 ml sterile water and centrifuging for 5 min at 3000 g. The pellet was then resuspended in 1 ml of sterile water and transferred into a 1.5 ml Eppendorf tube. Water was removed by centrifuging for 30 sec at 13000 g, and the pellet was resuspended again in 1 ml of water. From this, 100 µl (10^8^ cells) was pipetted into 1.5 ml Eppendorf tubes, one for each transformation, and centrifuged at 13000 g for 30 sec to remove the supernatant. The transformation mix was made by mixing 240 µl of PEG3350 (50% w/v), 36 µl of 1M LiAc, 50 µl single-stranded salmon sperm carrier DNA (2 mg ml^-1^), and 34 µL plasmid DNA (∼500 ng) in sterile water. The transformation mix was added to the cell pellet and mixed by vortexing vigorously. A transformation was done without plasmid as a control. Tubes were then incubated for 40 min at 42 °C. The supernatant was removed after centrifuging the cell suspension for 30 sec at 13000 g, and the pellet was resuspended in 200 µl YPD medium. Once delivered, the inoculum was spread on the SC-His plate for selection, and the plate was put in a 30 °C incubator for 3-4 days until colonies appeared. Transformation was confirmed by sanger sequencing of the mutated region of each plasmid. This was done by isolating genomic DNA of each yeast strain using the Zymoprep yeast plasmid miniprep II kit (Zymo Research) according to the manufacturer’s protocol and PCR amplifying the region of interest.

### Yeast drop test assay

Yeast cells were grown in YPD medium at 30 °C until they reached an OD_600_ of 1 or higher. Starting with the OD 1 culture, a 1:10 dilution series was prepared for each yeast strain. From each dilution (1, 1:10, 1:100, and 1:1000), 2 µl was spotted onto two YPD plates and each was incubated at 30 °C and 37 °C for 48-72 hours. Images were taken to compare the growth of each strain at 30 °C versus 37 °C.

### Yeast total RNA extraction with hot acidic phenol method

Yeast cells were grown overnight in 5 ml of YPD medium and pelleted by centrifugation at 1500 g for 3 min. Total RNA was extracted from the cell pellets using the hot acidic phenol method, as previously described ^72^. In brief, the cell pellets were resuspended in 1 ml of ice-cold water and transferred to a clean 1.5 ml Eppendorf tube. The cells were pelleted again by centrifugation at 1500 g for 3 min at 4 °C. The pellet was then resuspended in 400 µl of TES buffer (10 mM Tris-HCl pH = 7.5, 10 mM EDTA, 0.5% SDS), and 400 µl of preheated (65 °C) acid phenol was added. The mixture was vortexed vigorously for 10 sec and then incubated for 60 min at 65 °C with brief vortexing every 15 min for 10 sec. After 60 min, the tubes were placed on ice for 5 min and then centrifuged for 5 min at 21,000 g and 4 °C. The top aqueous layer was transferred to a clean 1.5 ml tube, and 400 µl of acidic phenol was added. The mixture was vortexed vigorously, kept on ice for 5 min, and then centrifuged for 5 min at 21,000 g and 4 °C. The aqueous layer was transferred to a clean tube containing 400 µl of chloroform, vortexed vigorously, and then centrifuged for 5 min at 21,000 g. Finally, the top aqueous layer, which contained the RNA, was transferred to a new tube. Ethanol precipitation was carried out by adding 40 ml of 3 M sodium acetate (pH = 5.3) and 1 ml of ice-cold 100% ethanol, incubating the mixture at -20 °C overnight, and then centrifuging for 30 min at 21,000 g and 4 °C. The pellet was washed with 70% ice-cold ethanol, dried, and resuspended in H_2_O.

### Ac^4^C analysis on yeast h45 site

Total RNA (5 µg) from each yeast transformant was treated with TURBO DNase (Invitrogen) by adding 1 µl of the enzyme, 5 µl of 10X buffer, H_2_O in a 50 µl total volume and incubating at 37 °C for 1 h. Then the RNA was isolated by using the Zymo RNA clean and concentrator-5 (Zymo Research) kit according to the manufacturer’s protocol.

Nucleotide resolution sequencing of ac^4^C in 18S rRNA-helix 45 was performed as previously described ^24^. RNA samples (1 µg) were treated with 100 mM NaCNBH_3_ or vehicle (H_2_O) in a final reaction volume of 100 μl. Reactions were initiated by addition of 1 M HCl to a final concentration of 100 mM and incubated for 20 min at room temperature. Reactions were stopped by neutralizing the pH by the addition of 30 μl 1 M Tris-HCl pH = 8.0. Then reaction volumes were adjusted to 200 μl with H_2_O, ethanol precipitated, and quantified by using a Nanodrop 2000 spectrophotometer. Yeast total RNA from individual reactions (200 pg) was incubated with specific h45 rev primer (4.0 pmol, TAATGATCCTTCCGCAGGTTCACCTAC) in a final volume of 20 μl. Individual reactions were heated to 65 °C for 5 min and transferred to ice for 3 min to facilitate annealing in SuperScript III reaction buffer (Invitrogen). After annealing, reverse transcriptions were performed by adding 5 mM DTT, 200 units SuperScript III RT, 500 μM dNTPs (use 250 μM dGTP) and incubating for 60 min at 55 °C. Reaction was quenched by increasing the temperature to 70 °C for 15 min and store at 4 °C. cDNA (2 μl) was used as template in 50 μl PCR reaction with Phusion Hot start flex (New England Biolabs). Reaction conditions: 1× supplied HF buffer, 2.5 pmol both forward (CGTCGCTAGTACCGATTGAATGG) and reverse primer (TAATGATCCTTCCGCAGGTTCACCTAC), 200 μM each dNTPs, 2 units Phusion hot start enzyme, 2 μl cDNA template. PCR products were run on a 2% agarose gel, stained with SYBR safe, and visualized on UV transilluminator at 302 nm. Bands of the desired size were excised from the gel and DNA extracted using QIA-quick gel extraction kit (Qiagen) and submitted for Sanger sequencing (Azenta) using the forward PCR primer. Processed sequencing traces were viewed using 4Peaks software. Peak height for each base was measured and the percent misincorporation was determined using the equation: Percent misincorporation = (Peak intensity of T) / (Sum of C and T base peaks) × 100%.

### Human cell line culture and etoposide induction

IMR90 (Primary human lung fibroblast) and HEK293T (Human kidney cell line) were described previously. IMR90 and HEK293T cells were cultured in DMEM supplemented with 10% fetal bovine serum (FBS) and 100 units ml^−1^ penicillin and 100 µg ml^−1^ streptomycin (Invitrogen). IMR90 cells were cultured under physiological oxygen (3%). IMR90 cells with a population doubling of less than 40 were used for experiments. For etoposide-induced senescence, IMR90 cells were transiently treated with 100uM of etoposide (Sigma-Aldrich Catalog# E1383) for 48 hours. Cells were replaced with untreated media and assayed at designated time points.

### Immunoblot

IMR90 cells were lysed in buffer containing 20 mM Tris pH = 7.5, 137 mM NaCl, 1 mM MgCl_2_, 1 mM CaCl_2_, 1% NP-40, 10% glycerol and supplemented with Halt Protease Inhibitor Cocktail (ThermoFisher #78430) and Benzonase (MilliporeSigma #70746-3). Cells were incubated at 4 °C shaking for at least 1 hour. Sodium dodecyl sulfate (SDS) was added to a final concentration of 1% and boiled for 10 minutes. Samples were centrifuged at 13000 RPM and supernatant (lysate) was quantified for protein concentration. Lysates were subjected to electrophoresis and transferred to a 0.2 μm nitrocellulose membrane. The membrane was blocked with 5% Milk for at least 30 minutes shaking and rinsed 3 times using Tris-buffered saline (TBS) supplemented with 0.1% Tween (TBS-T). The membrane was incubated overnight at 4 °C with primary antibodies (See **Key Resource Table S3**) in TBS-T in 5% Bovine Serum Albumin (BSA) and probed with horseradish peroxidase-conjugated secondary antibodies and developed with SuperSignal West Pico Plus chemiluminescent substrate (ThermoFisher #34580) and imaged using an Amersham Imager 600.

### Human total RNA Isolation and quantitative PCR

The harvested cells were washed and incubated with 1 ml of TRIzol reagent and incubated for 5 minutes. 200 μl of chloroform was added per sample and vortexed. Samples were centrifuged and clear supernatant was harvested in new Eppendorf tube containing 318 μl of 100% ethanol. Samples were transferred to the RNeasy column (Qiagen #74106) and centrifuged. The columns were washed with 500 μl of RW1 buffer and treated with DNase (Qiagen #79254) for 30 minutes at room temperature. The columns were then washed with 500 μl of RW1 buffer followed by 500 μl of RPE buffer and eluted with 30 μl of RNase free water. Isolated RNA was reversely transcribed using High-Capacity cDNA Reverse Transcription Kit (ThermoFisher #4368813). cDNA was diluted 1:10 in H_2_O and was used for quantitative PCR using Power SYBR Green PCR Master Mix (ThermoFisher #4368708) following manufacturer protocol and run on Applied Biosystem QuantStudio Real-Time PCR System using targeted primer and reference primer sets (See **Key Resource Table S4**).

## SUPPLEMENTAL FIGURES

**Supplemental Figure 1.**
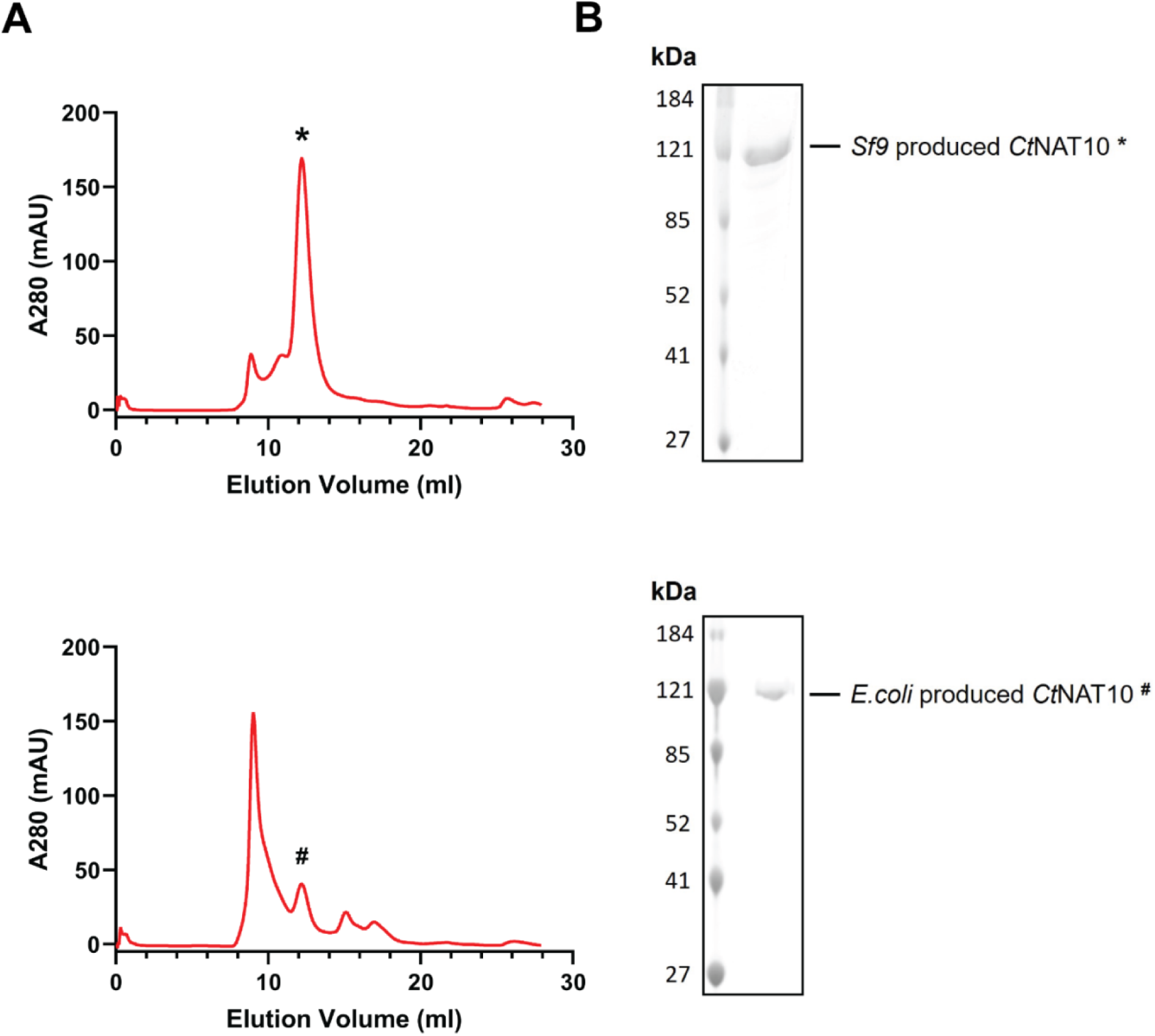
Production and purification of recombinant *Ct*NAT10 from *Sf9* and *E.Coli*. **(A)** Superdex-200 size-exclusion chromatogram of dimeric N-terminally His-tagged *Ct*NAT10 purified from *Sf9* (*) and *E.Coli* (^#^). **(B)** SDS-PAGE (12.5% acrylamide) of intact, homogeneous *Ct*NAT10 protein produced in *Sf9* (*) and *E.Coli* (^#^). Polypeptides were visualized by staining with Coomassie Blue dye. The positions and sizes (in kDa) of marker proteins are indicated on the left.

**Supplemental Figure 2.**
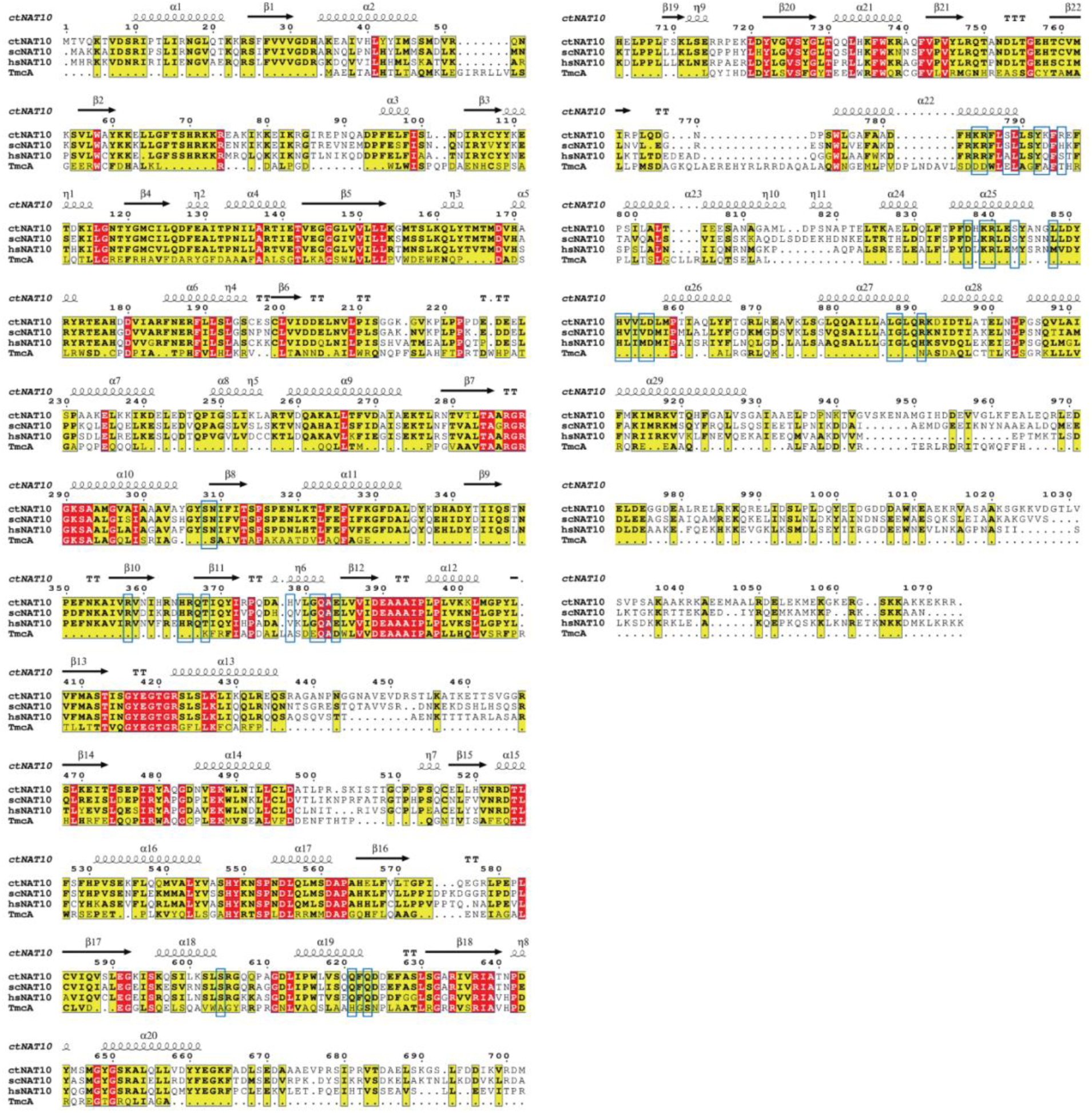
Sequence alignment of NAT10 homologs. The aligned species are *Chaetomium thermophilum* (ct), *Saccharomyces cerevisiae* (sc), *Homo sapiens* (hs) and *Escherichia coli* (TmcA). Secondary structure elements are indicated for the *E.coli* produced recombinant *Ct*NAT10/cytidine-amide-CoA/ADP cryo-EM structure. Conserved amino acids are in white in red-filled rectangles. Similar residues are in bold in yellow-filled rectangles, surrounded by black lines. Residues that are involved in *Ct*NAT10 dimerization surface are boxed in blue rectangles. Alignment was performed using ESPript 3 ^1^.

**Supplemental Figure 3.**
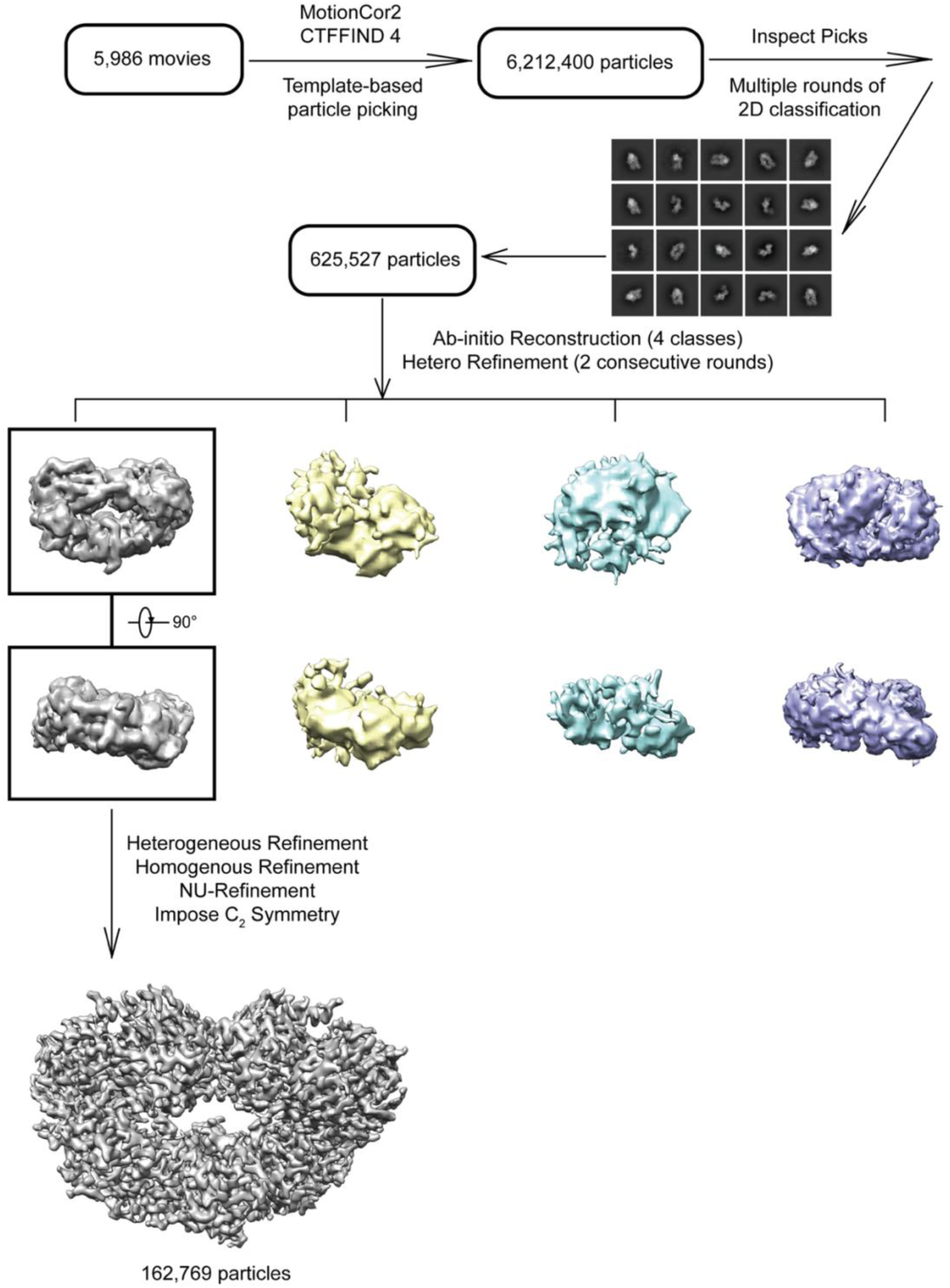
Cryo-EM image processing workflow of *Ct*NAT10 structures. Workflow, 2D and 3D classification scheme for *Ct*NAT10_Sf9_ /cytidine-amide-CoA/ADP is shown. The workflows for the other two structures are similar.

**Supplemental Figure 4.**
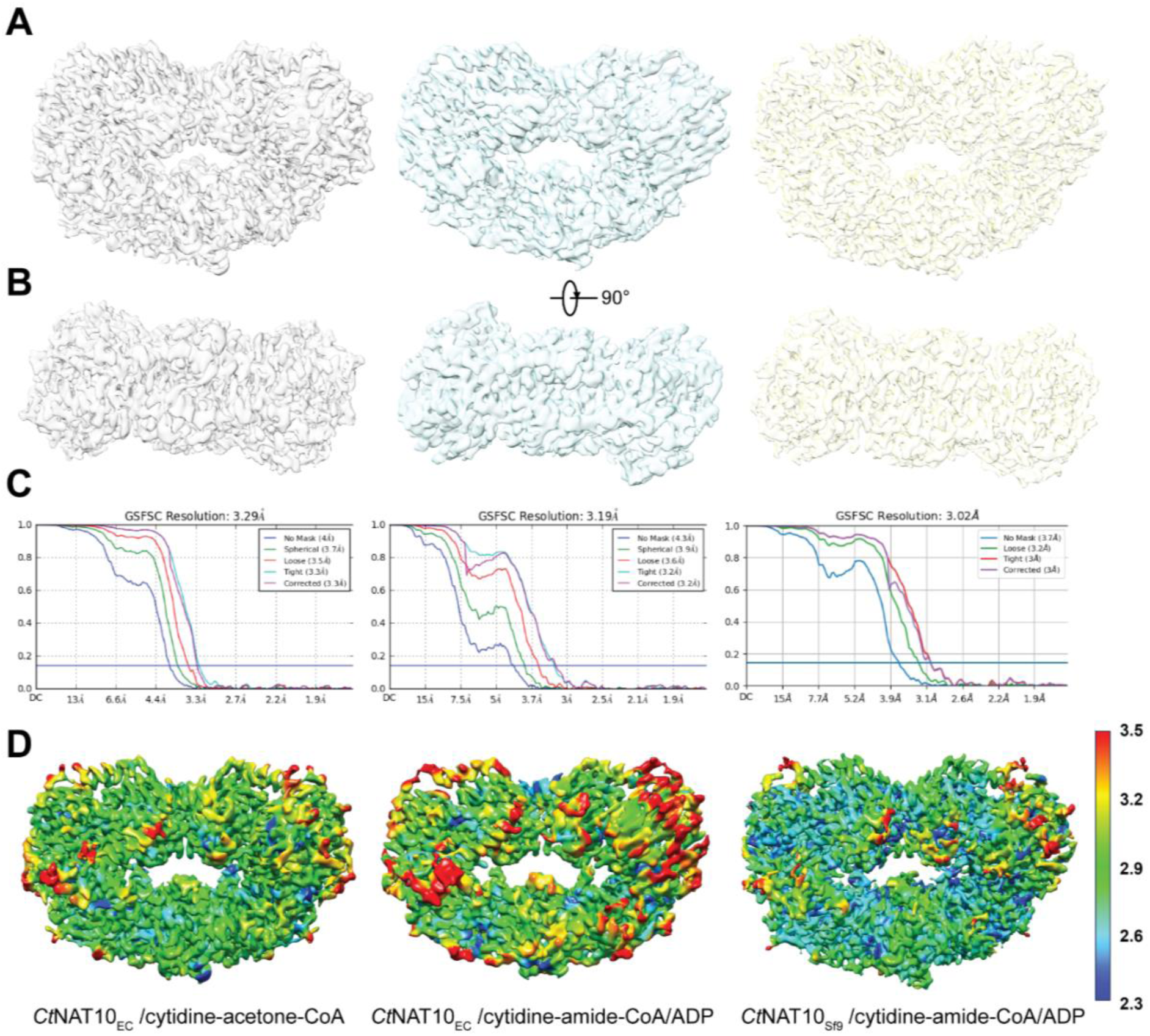
Cryo-EM maps quality control and resolution estimation. **(A)** Front and **(B)** bottom view, **(C)** Fourier shell correlation (FSC) curves and **(D)** local resolution estimation for the cryo-EM density of *E.coli* produced recombinant *Ct*NAT10/cytidine-acetone-CoA, *E.coli* produced recombinant *Ct*NAT10/cytidine-amide-CoA/ADP and *Sf9* produced recombinant *Ct*NAT10/cytidine-amide-CoA/ADP (from left to right), respectively. The volume threshold levels for displaying the three Cryo-EM maps are 0.264, 0.229 and 0.394, respectively. All FSC curve plots are generated by cryoSPARC.

**Supplemental Figure 5.**
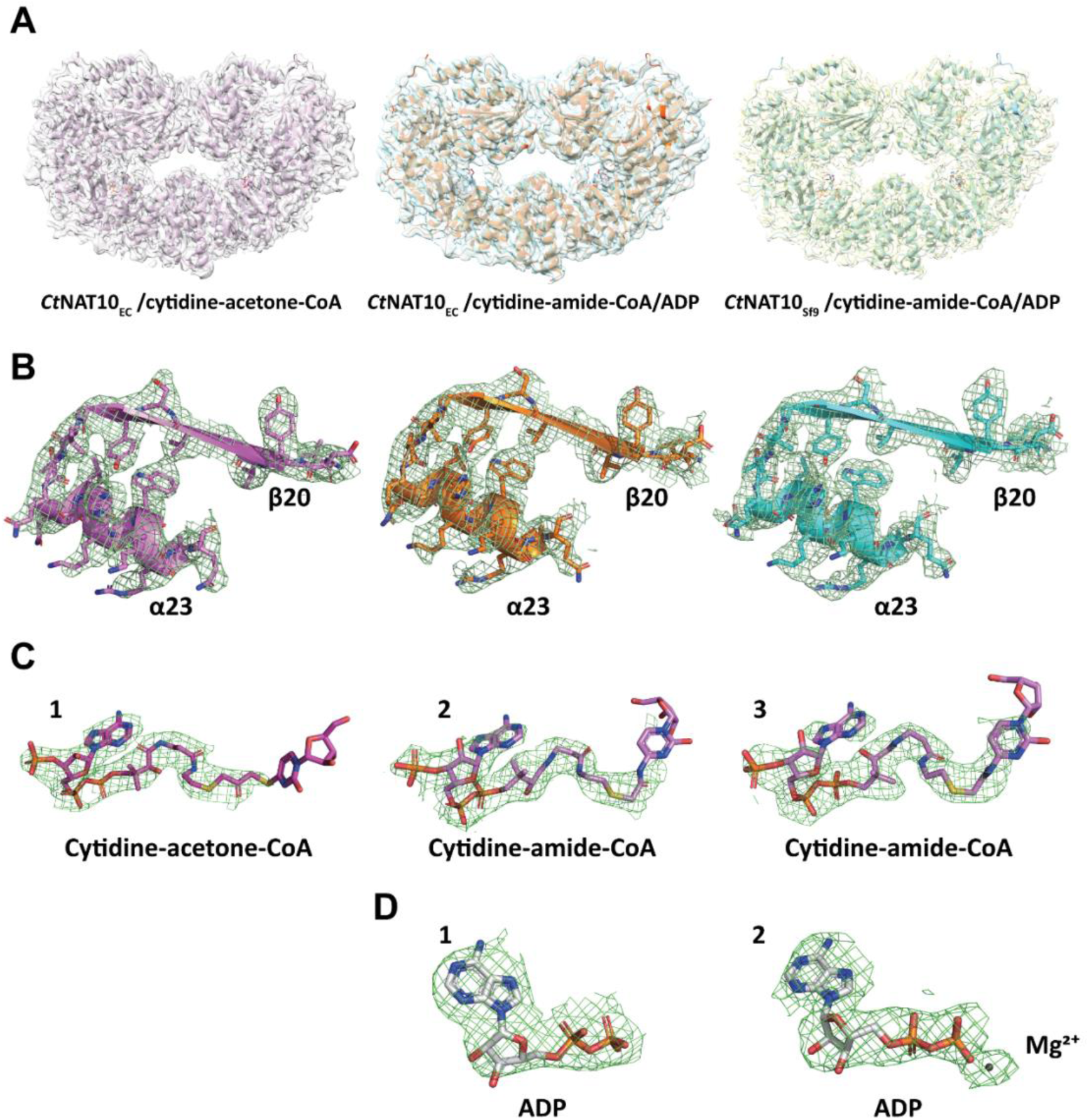
Atomic models fitted into the cryo-EM maps. **(A)** Atomic models of *E.coli* produced recombinant *Ct*NAT10/cytidine-acetone-CoA (lavender), *E.coli* produced recombinant *Ct*NAT10/cytidine-amide-CoA/ADP (orange) and *Sf9* produced recombinant *Ct*NAT10/cytidine-amide-CoA/ADP (cyan) cryo-EM structures, were fitted into the corresponding cryo-EM maps using Chimera, respectively. **(B)** The “fit-in-map” of the catalytically important secondary structure elements β20-α23 (residue 720-741) for all three *Ct*NAT10 structures are shown, for demonstrating the cryo-EM map quality in detail. A contour level of 8 σ was applied to all the maps. **(C) and (D)** Cryo-EM density of ligands. Density of cytidine-CoA conjugates (C - 1, 2, 3) and ADP- (Mg^2+^) (D - 1, 2), for three *Ct*NAT10 structures (the ligands are in the same column with the *Ct*NAT10 structure that they are bound to). A contour level of 7 σ is shown in C-1, a contour level of 10 σ is shown in D-1 and a contour level of 8 σ is shown in C-2, C-3 and D-2. All mesh of cryo-EM maps and corresponding fit-in model figures are depicted using Pymol.

**Supplemental Figure 6.**
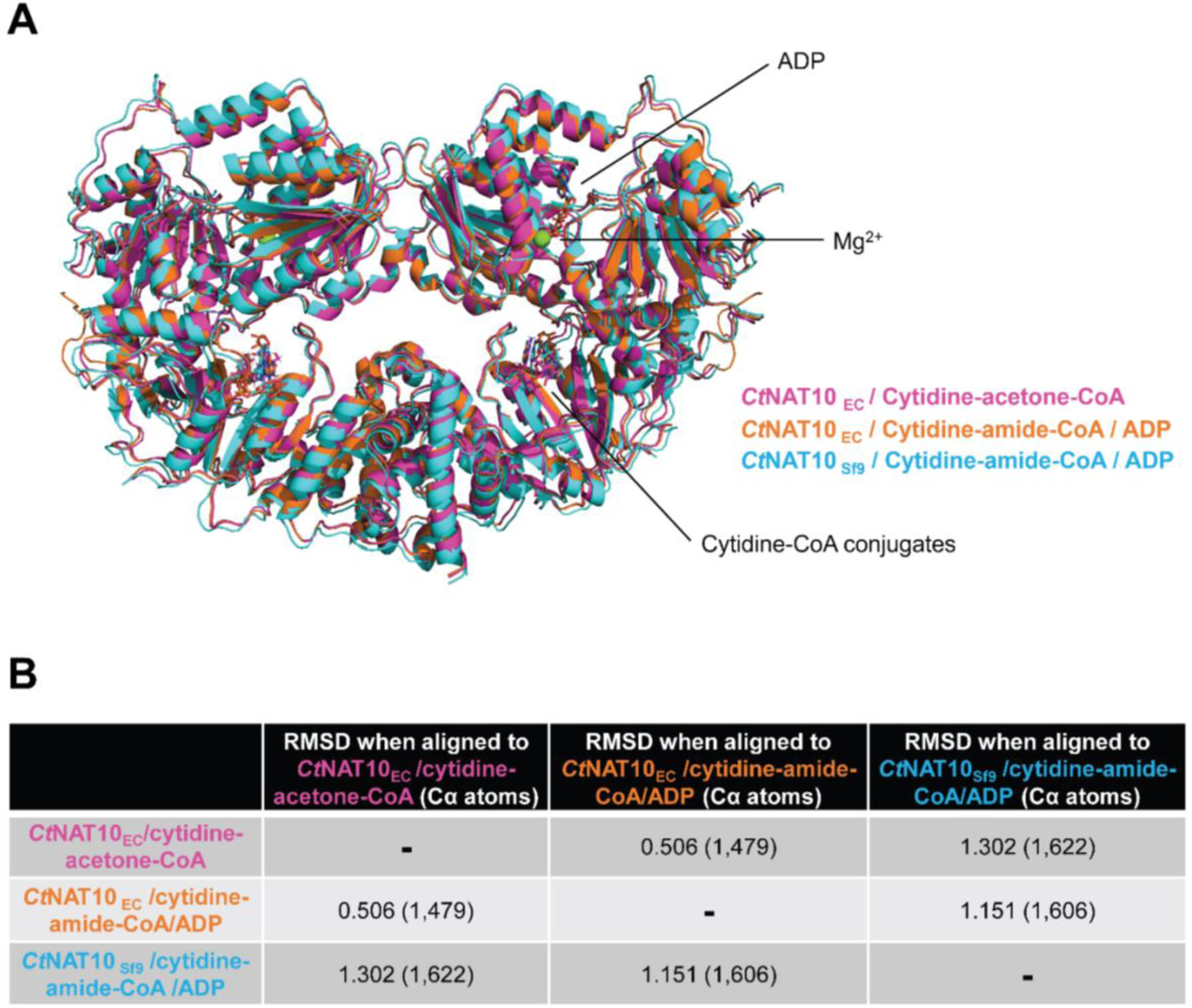
Comparison of three *Ct*NAT10 cryo-EM structures. **(A)** Superimposition of the *E.coli* produced recombinant *Ct*NAT10/cytidine-acetone-CoA (magenta), *E.coli* produced recombinant *Ct*NAT10/cytidine-amide-CoA/ADP (orange) and *Sf9* produced recombinant *Ct*NAT10/cytidine-amide-CoA/ADP (cyan) cryo-EM structures. **(B)** Root-mean-squared deviations of the three *Ct*NAT10 structures.

**Supplemental Figure 7.**
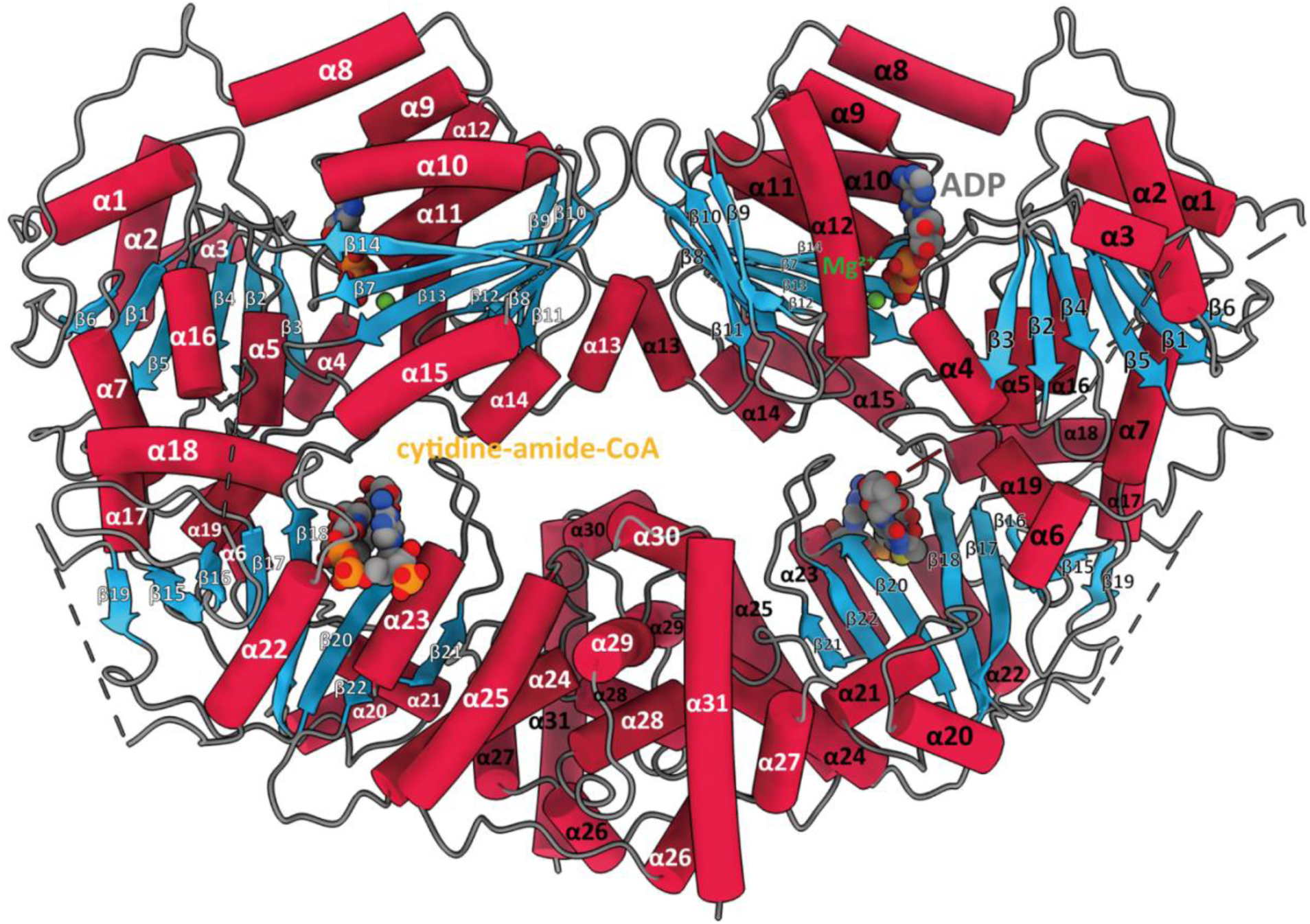
Cartoon representation of *Ct*NAT10 structure with labelled secondary structures. Since the three *Ct*NAT10 structures reported in this paper superimposed well, here we use *Sf9* produced recombinant *Ct*NAT10/cytidine-amide-CoA/ADP structure as the representative. α-helix (crimson) and β-sheet (skyblue) are labelled and numbered in the order from N-terminus to C-terminus. One of the subunits of the two-fold symmetric *Ct*NAT10 dimer is labeled with black, whereas the other is labeled with white to facilitate visualization of the dimeric assembly.

**Supplemental Figure 8.**
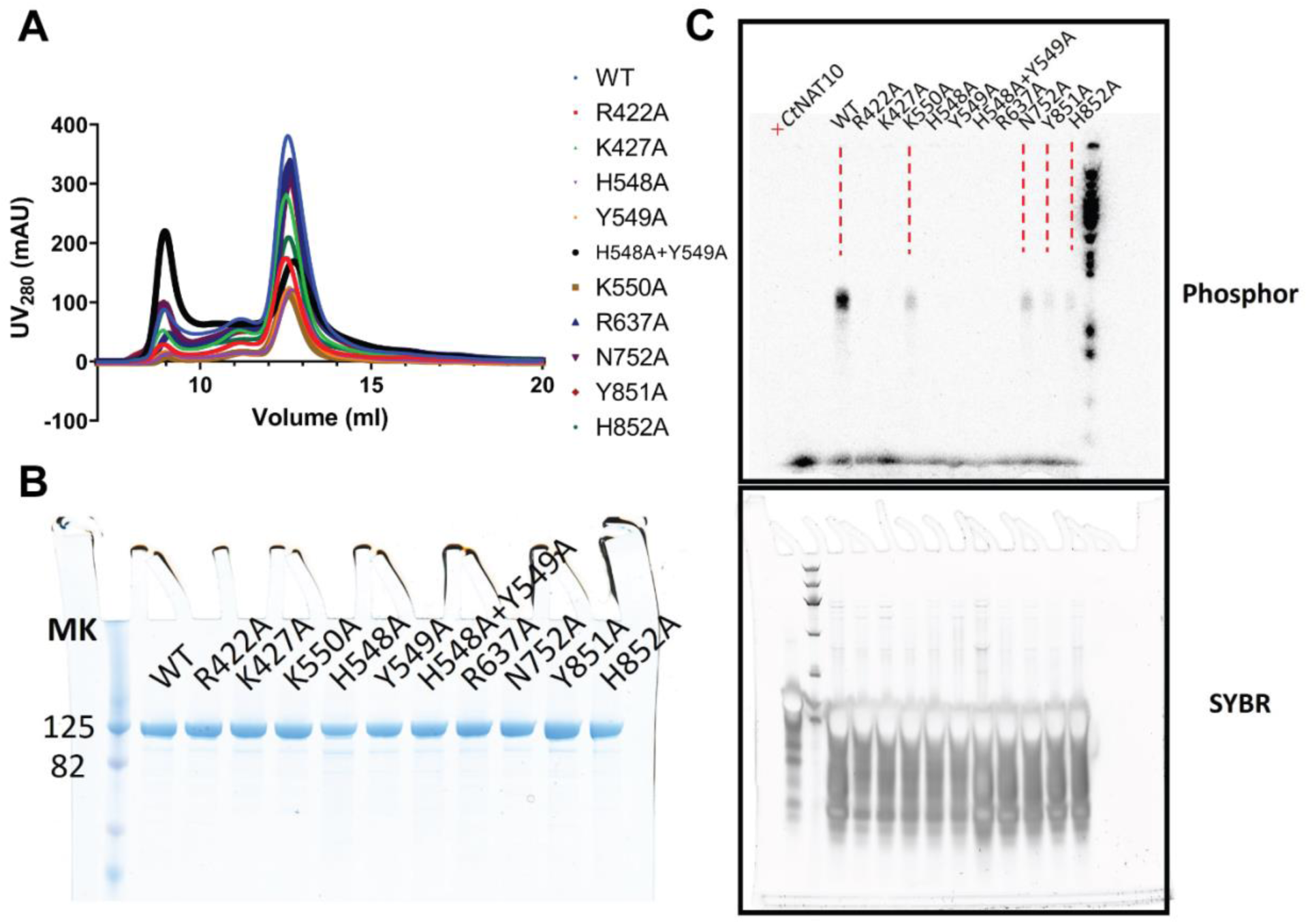
Purification and activity assessment of recombinant *Ct*NAT10 single-alanine mutant proteins. **(A)** Superdex-200 size-exclusion chromatogram of dimeric N-terminally His-tagged *Ct*NAT10 alanine mutants purified from Sf9. **(B)** SDS-PAGE (12.5% acrylamide) of intact, homogeneous *Ct*NAT10 alanine mutant proteins produced in Sf9. Polypeptides were visualized by staining with Coomassie Blue dye. The positions and sizes (in kDa) of marker proteins are indicated on the left. **(C)** Representative autoradiogram of the gel for evaluating the effect of the indicated mutant constructs on *Ct*NAT10 *in vitro* acetyltransferase activity towards a h45 rRNA fragment. The assays were performed essentially as described in the time course assay shown in Figure 1A. The sample on the very left lane represent no protein negative control.

**Supplemental Figure 9.**
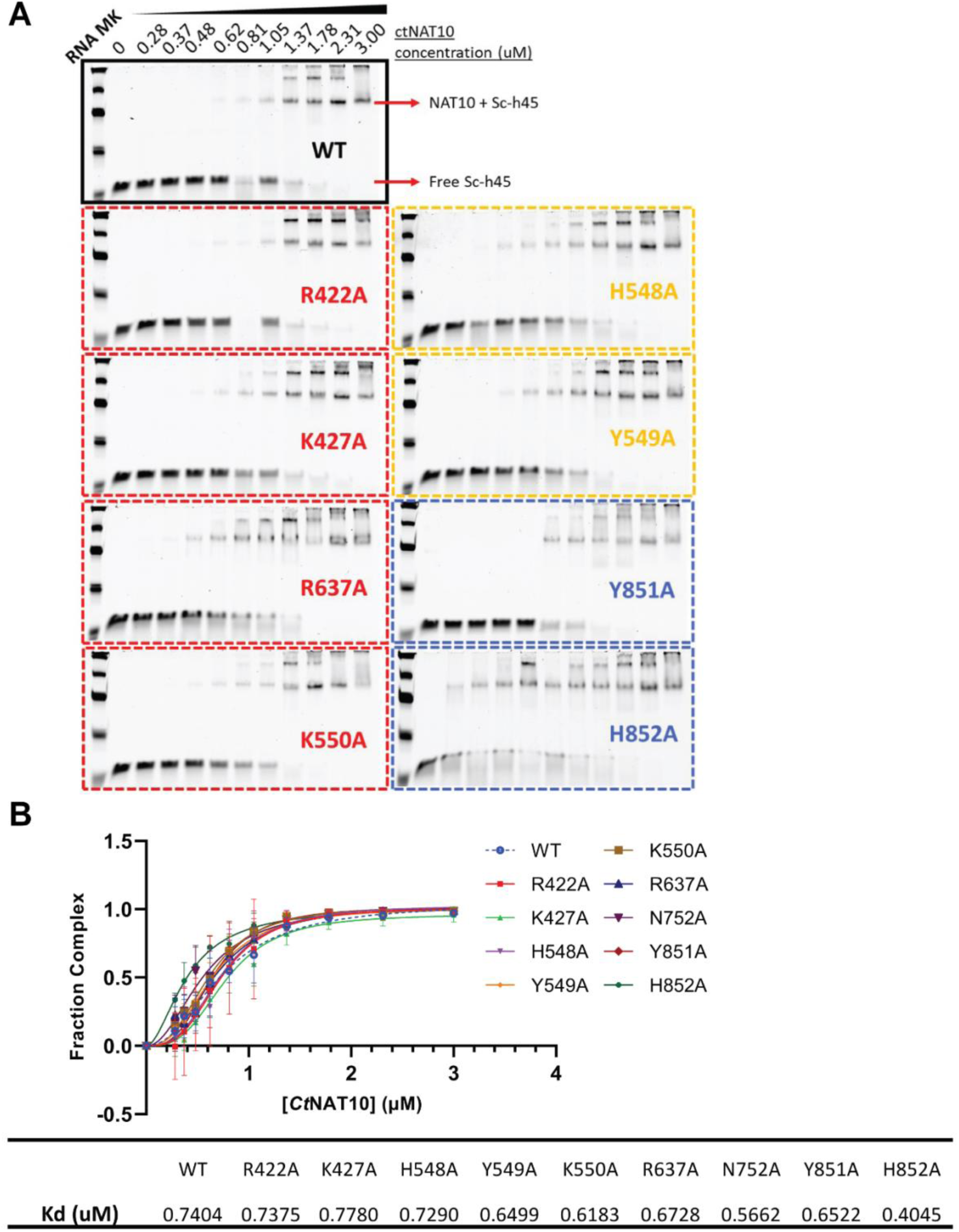
EMSA of purified *Ct*NAT10 mutants with h45 rRNA substrate binding. **(A)** 6% polyacrylamide native gels were stained with SYBR™ Gold to visualize RNA fragment shift. RNA input concentration fixed at 0.25 µM for all the samples, which were premixed and run with various *Ct*NAT10 alanine-mutant concentrations (0-3.0 µM, number indicated in figure). The color-coding of the mutant construct names correspond to their locations defined in Figure 4D. RNA-*Ct*NAT10 complex bands and free RNA probe bands are indicated with arrows. **(B)** Fraction complex value derived from the EMSA data, versus *Ct*NAT10 concentration was fitted using the Hill equation (assorted-colored lines). Quantification of “fraction free-RNA” (F_RNA_) was carried out using ImageJ to measure the intensity of the free RNA band, then normalized against the intensity of [protein] = 0 µM. The value of fraction complex = 1 - F_RNA_. Averaged values of three technical replicates with S.D. values are shown for each data point and their corresponding error bars. The mean K_d_ value for all *Ct*NAT10 mutants complex from three repeat experiments was estimated and listed in the table at the bottom.

**Supplemental Figure 10.**
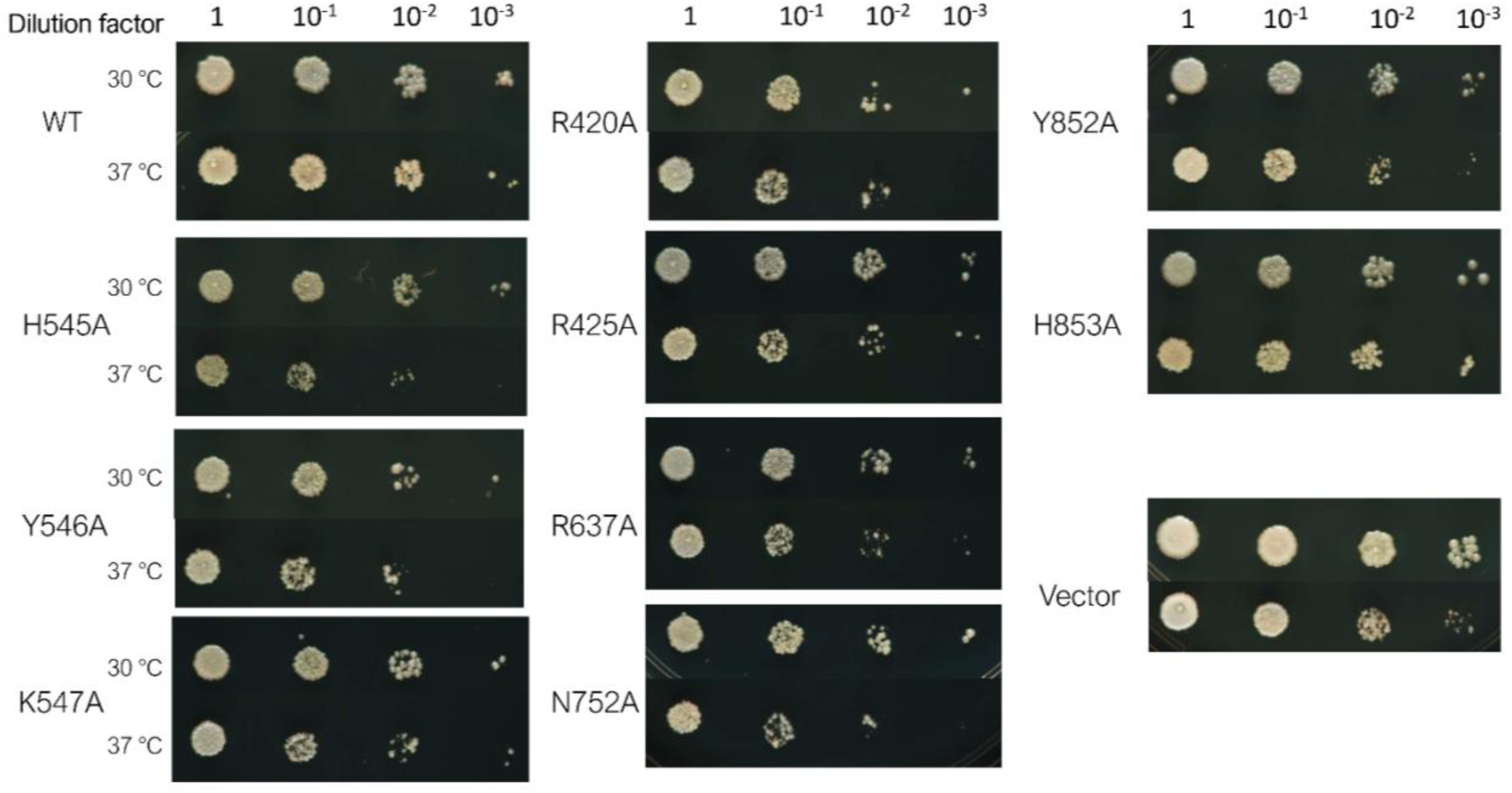
Growth assay data for Figure 5. Strains were grown, diluted, plated on YPD plates, and grown at the indicated temperatures for 48 h.

**Supplementary Figure 11.**
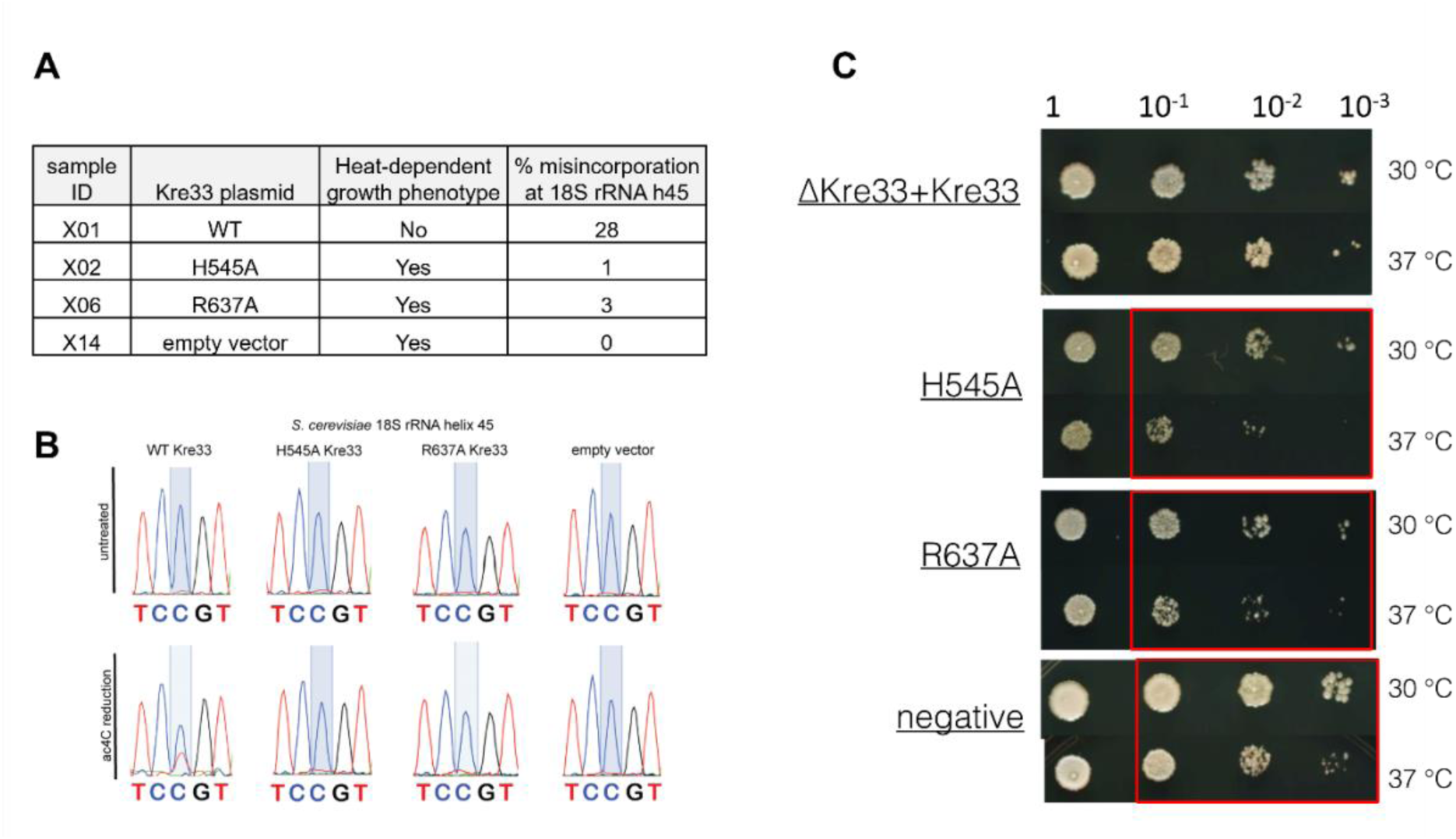
Rescue of high-temperature growth defect correlates with rescue of RNA cytidine acetylation. **(A)** Strains were transformed as described in Figure 5A and total RNA was isolated by hot acidic phenol method. The presence of ac4C at 18S SSU helix 45 was then assessed using a quantitative ac4C sequencing reaction described in detail elsewhere ^2^. *Sc*Kre33 mutants that successfully rescue exhibit no heat-dependent growth phenotype and also display rescue of 18S SSU helix 45 acetylation (WT only). **(B)** Exemplary ac4C sequencing data. **(C)** *Sc*Kre33 mutants that do not rescue exhibit a reduced growth at higher temperatures and do not display rescue of 18S SSU helix 45 acetylation (H545A, R637A, empty vector) are marked with red boxes.

